# Design and optimization of novel succinate dehydrogenase inhibitors against agricultural fungi based on Transformer model

**DOI:** 10.1101/2024.02.20.581130

**Authors:** Yuan Zhang, Jianqi Chai, Ling Li, Wenqian Zhao, Yuanyuan Chen, Liangyun Zhang, Zhihui Xu, Chunlong Yang, Cong Pian

**Author notes:** These authors are corresponding authors: Cong Pian, Chunlong Yang, Zhihui Xu. These authors contributed equally: Yuan Zhang, Jianqi Chai.

## Abstract

Succinate dehydrogenase inhibitors (SDHIs) are a promising class of fungicides targeting the energy production pathway of pathogenic fungi. However, overuse has led to resistance, necessitating the development of new and effective SDHIs. This study takes the Transformer model to generate a customized virtual library of potential SDHIs. These candidates were then meticulously screened based on expert knowledge and synthetic feasibility, ultimately yielding several pyrazole carboxamide derivatives as the promising leads. Subsequent synthesis, antifungal activity testing, and structural optimization further refined these leads into potent SDHI candidates. This work marks the first application of a generative model to SDHI design, establishing a robust workflow for virtual library generation, screening, activity evaluation, and structure optimization. This provides one way for the rational design of future SDHIs, not only against fungi, but potentially other agricultural pathogens as well.

## Introduction

Succinate dehydrogenase is an important biological enzyme in the mitochondrial oxidative phosphorylation process. It is also an essential enzyme in the tricarboxylic acid cycle and mitochondrial electron transfer. It consists of four subunits: SDHA, SDHB, SDHC and SDHD [1]. Succinate dehydrogenase inhibitors (SDHIs) are a type of commercial fungicides that are very important in agriculture. This type of fungicides has become the next most popular fungicide after quinone outside respiration inhibitors and ergosterol biosynthesis inhibitor fungicides. SDHIs can block the mitochondrial tricarboxylic acid cycle and respiratory chain electron transport [2], thereby exerting its fungicidal activity by inhibiting the activity of succinate dehydrogenase [3]. As of now, there were 24 SDHIs on the fungicide resistance action committee [4-5]. However, the frequent use of SDHIs, their single site of action, prolonged drug use, and increased drug dosage have resulted in many pathogenic fungi developing serious resistance to existing SDHIs [6]. Therefore, it is urgent to develop new SDHIs with unique molecular structures.

In recent years, deep learning technology has continued to develop, and various deep generative models have been proposed. Deep generative models have achieved a series of breakthroughs in the field of drug discovery. Recurrent neural network (RNN) is a type of model commonly used to capture sequence information. This type of model is very effective when processing data with sequential or temporal information. Since the simplified molecular input line entry system (SMILES) representation of molecules reflects the information of atoms and bonds of molecules with sequence information [7]. Molecules can be generated based on RNN models [8-10]. The generative model based on variational autoencoder (VAE) consists of two networks: (1) encoder, which maps the input into a low-dimensional latent vector; (2) decoder, which maps the latent vector into the newly generated data [11]. Both the encoder and decoder can be common RNN or convolutional neural networks (CNN) models. Therefore, VAE can be used to extract SMILES information of molecules and can also be used to extract molecular graph information [12-14]. The generative model based on generative adversarial network (GAN) is composed of a generator and a discriminator [15]. The generator aims to generate data similar to the training data, and the discriminator is a classification model designed to distinguish whether the input data is real data or the data generated by the generator. The generator and discriminator are trained adversarially until the discriminator cannot distinguish between real data and data generated by the generator, which means that the generator is able to generate reasonable molecules. Both the generator and the discriminator can be common RNN or CNN models. Therefore, the generative adversarial networks can be used to extract SMILES information of molecules, and can also be used to extract molecular graph information [16-19]. The generative model based on Transformer [20] uses a self-attention mechanism to capture the correlation between data and uses an autoregressive form for molecule generation [21-23]. Most of the above models are based on the generation of single atom. Some researchers have also begun to study molecular generation models based on chemical molecular scaffold information [24-26], which lays the foundation for the subsequent generation of active molecules with new scaffolds. In addition to using the information of the molecule itself for ligand-based drug design, the three-dimensional structure information of the molecular target protein is also used in the molecular generation models to achieve structure-based drug design [27-29]. Up to now, many researchers have used the models proposed above to design specific chemical small molecules. William J. Godinez et al. established a JAEGER model based on junction-tree VAE, designed compounds that inhibit malaria, synthesized and analyzed two screened compounds, and found that both compounds exhibited powerful anti-malarial activity [30]. Lijuan Yang et al. used a Transformer-based generative model to design novel BRAF inhibitors, and demonstrated through molecular docking and binding mode analysis that their generative model can generate high-quality BRAF inhibitor candidate compounds [31]. Yueshan Li et al. used a CRNN-based generative model to design a customized virtual compound library targeting RIPK1 and performed virtual screening to identify an effective and selective RIPK1 inhibitor [32]. Moret et al. used chemical language models to design PI3Kγ inhibitors in the form of pre-training and fine-tuning. The generated molecules and several derivatives were synthesized and tested for activity. Finally, they found that these compounds effectively inhibited PI3Kγ activity [33]. The above studies show that deep generative models have shown promise in the design of inhibitors for targets related to human diseases, but there has been no relevant exploration in the field of agricultural fungicides. In this study, a framework combining artificial intelligence technology for succinate dehydrogenase inhibitors design was proposed, in which a customized virtual compound library for SDHIs was designed using the Transformer model, which contained a total of 91,961 molecules. After a series of molecular docking analyses and manual screening, some compounds were finally selected for subsequent synthesis and bioactivity testing, and discovered a compound has appreciable inhibitory activity against phytopathogenic fungi. We conducted two rounds of optimization on the structure of the compound with the best inhibitory effect on tested fungi, and finally obtained some potential SDHIs with obvious antifungal activity, laying the foundation for the design of new succinate dehydrogenase inhibitors. To the best of our knowledge, this article is the first time that the Transformer model as a deep generative model is applied to the design of SDHIs, opening up a new path for the development of new agricultural fungicides.

## Results

Fig. 1 shows the work process of this article. The data collection section shows the source of the data. The data of this article are from documents provided by the fungicide resistance action committee in 2020. Based on this document, commercialized succinate dehydrogenase inhibitors were collected. The process section shows the workflow of this article, using the data of the MOSES benchmark platform to pre-train the model, using SDHIs to fine-tune the model, generating molecules, screening, molecule docking, synthesizing and evaluating antifungal activity. The generation model section shows the model, and the specific implementation can be found in the methods. In this article, the trained model was used to generate 100,000 molecules. After removing invalid molecules and duplications, there are a total of 91,961 compounds. These compounds were further screened and analyzed.

**Fig. 1.**
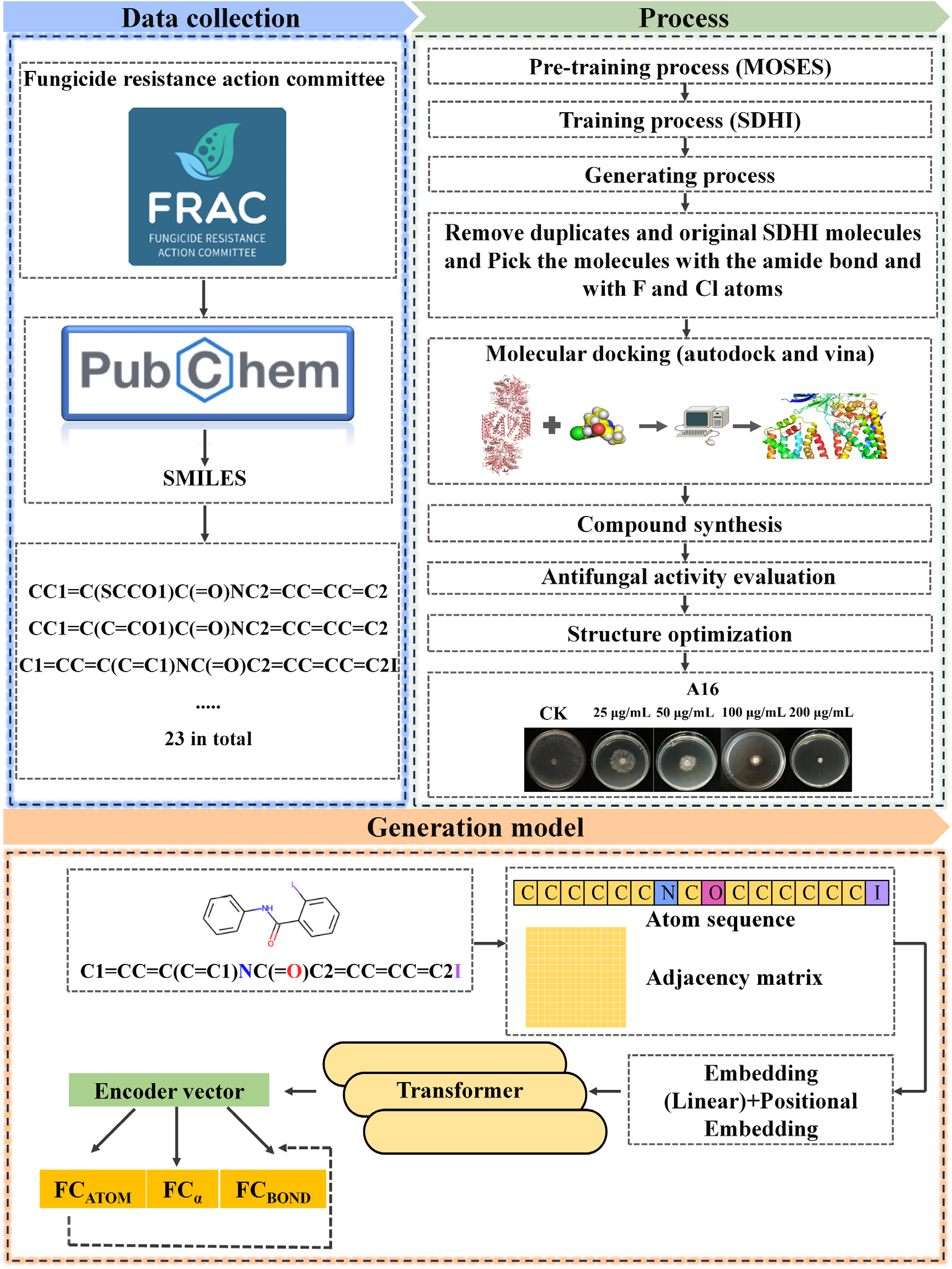
Process for designing, screening and optimizing succinate dehydrogenase inhibitors against agricultural fungi based on the Transformer model. The Data collection part describes the data collection process, the process part describes the entire process, and the generation model part briefly describes the implementation of the generated model.

### Validity, novelty and uniqueness analysis

To evaluate the ability of the model to generate molecules, the validity, uniqueness, and novelty of the generated molecules were calculated. As shown in Fig. 2A, the validity of the molecules generated by the model is 0.92567, the uniqueness is 0.993, and the novelty is 1. Higher validity indicates that well-defined chemical constraints are captured, higher uniqueness indicates that the model does not collapse to produce only a few typical molecules, and high novelty indicates that the model can generate more new molecules. The higher novelty and uniqueness laid the foundation for the subsequent screening of several active compounds.

**Fig. 2.**
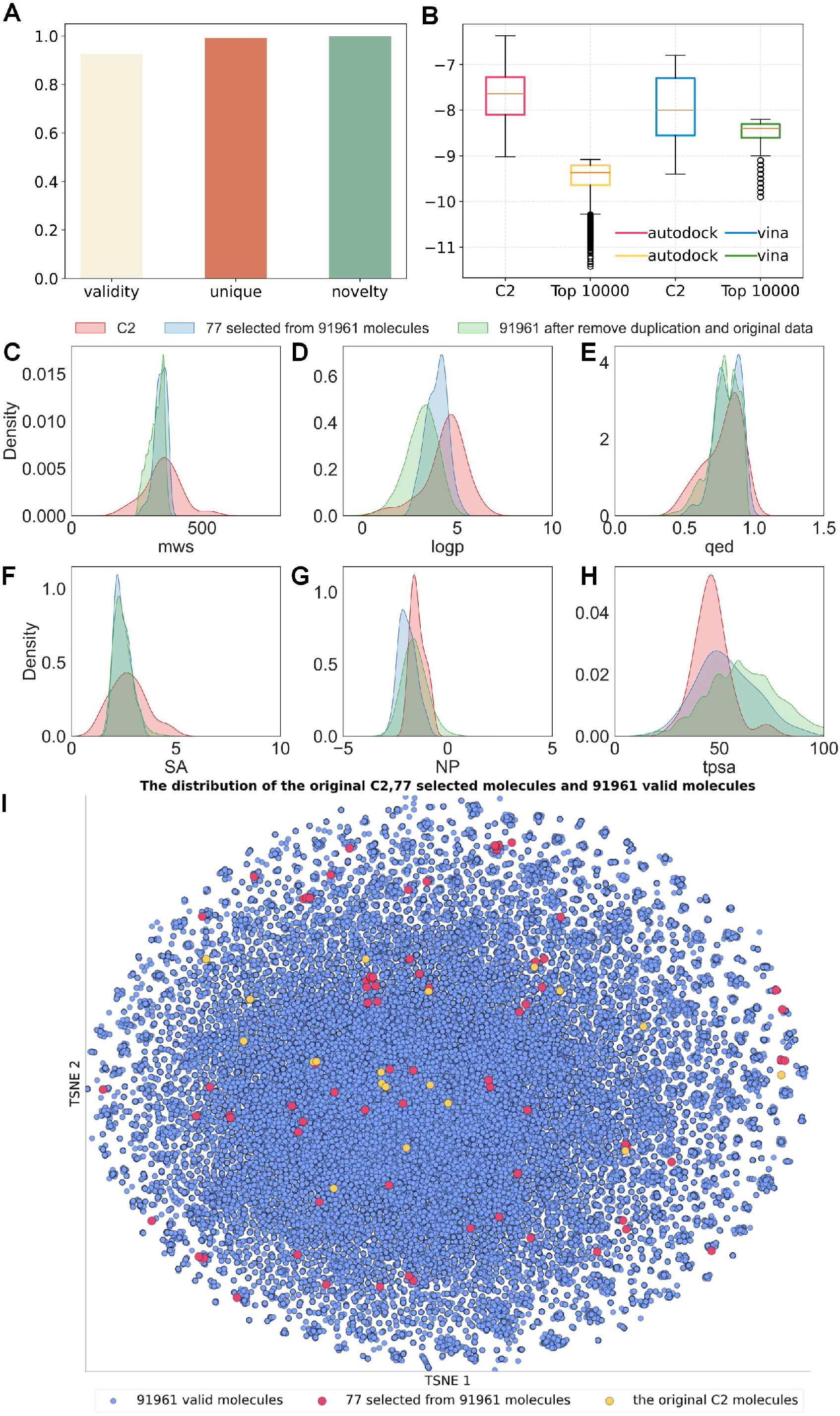
Results of model evaluation, docking scores, and molecular property distribution of generated molecules. Figure 2A shows the validity, uniqueness, and novelty of the model. Figure 2B shows the distribution of free energy. Figure 2C-2H diagram show the molecular weight, logp, drug-likeness, synthetic accessibility, natural product-likeness, and relative topological surface area distribution of SDHIs and generated molecules. Figure 2I shows the distribution of SDHIs and generated molecules in two-dimensional space.

### Molecular properties of generative molecules and SDHIs

In order to compare the physicochemical properties of generated molecules with SDHIs, some physicochemical properties of SDHIs, 77 selected generated molecules, and 91961 unselected generated molecules were calculated. Fig. 2C-H respectively show the distribution of the molecular weight, logp, quantitative estimate of drug-likeness (QED), synthetic accessibility (SA), natural product-likeness (NP) and relative topological surface area (TPSA) of the above molecules. The red part belong to SDHIs, the blue part belong to 77 selected molecules, and the green part belong to 91961 unselected molecules. It can be seen that the molecular weights of SDHIs are mostly concentrated between 300 and 400, and most of the 77 molecules selected from generated molecules and 91961 unselected generated molecules are also in this range inside. Most of the logp of SDHIs are concentrated between 3 and 5, while most of the 77 selected molecules and 91961 unselected generated molecules are concentrated between 2 and 4. This suggests that the generated molecules are more hydrophilic. Most of the QED of SDHIs, 77 selected generated molecules and 91,961 unselected generated molecules are concentrated between 0.5 and 1. Most of the of the SA of SDHIs, the 77 selected generated molecules and the unselected 91,961 generated molecules are mostly concentrated between 2 and 3. But the SA distribution shape of SDHIs are similar to the 77 selected generated molecules. Most of the NP of SDHIs, the 77 selected generated molecules and the unselected 91,961 generated molecules are concentrated between -2 and 0. In terms of TPSA, most of the generated molecules are higher than the relative topological surface area of SDHIs, which indicates that the generated molecules are more soluble in water and can pass through the cell membrane more easily.

### Distribution of generated molecules and SDHIs

To visualize the spatial distribution of generated molecules and SDHIs, the distribution of SDHIs, 77 selected generated molecules, and 91961 unselected generated molecules in two dimensions were visualized using t-SNE. As shown in Fig. 2I, the distribution of yellow points represents the distribution of SDHIs in two-dimensional space, the distribution of red points represents the distribution of 77 selected generated molecules in two-dimensional space, and the distribution of blue points represents the distribution of 91961 unselected generated molecules in two-dimensional space. It can be seen that the distribution of the generated molecules covers the entire chemical space of SDHIs in two-dimensional space.

### Similarity analysis of generated molecules and SDHIs

To compare the similarities of the generated molecules to SDHIs, the similarities between SDHIs and each of 77 selected generated molecules and 91,961 unselected generated molecules were calculated. The results are shown in supplementary Fig. 1 and supplementary Fig. 2. The green curve is the similarity between the SDHIs and the 77 selected generated molecules, and the red curve is the similarity between the SDHIs and 91,961 unselected generated molecules. Most similarities between our generated molecules and SDHIs are clustered between 0.2 and 0.6, which means the high novelty of the generated molecules.

### Results of molecular docking

The molecular docking of chicken heart succinate dehydrogenase (PDB:2FBW) and each molecule of the SDHIs was performed. And the molecular docking of chicken heart succinate dehydrogenase (PDB:2FBW) and each molecule of the 91961 generated molecules was performed too. The docking scores of 91961 generated molecules were ordered from small to large, taking the free energy of the top 10000 molecules. The distribution of docking scores for SDHIs and the top 10000 molecules generated by the generative model are shown in Fig. 2B. It can be found that molecules generated by the generative model had lower docking scores than SDHIs, suggesting that these molecules may have better activity. It can be known from previous studies that the oxygen atom in the amide bond of SDHIs often forms hydrogen bonds with B_W173 and D_Y91 [34-35]. The structures and docked conformational images of three molecules selected from the generated molecules for antifungal activity evaluation are shown in Fig. 3. It can be seen from the conformation that all the three selected molecules form hydrogen bonds with TRP-173 and TYR-58 in both of the autodock-docking conformation and the vina-docking conformation. This shows that the model can learn some properties of SDHIs.

**Fig. 3.**
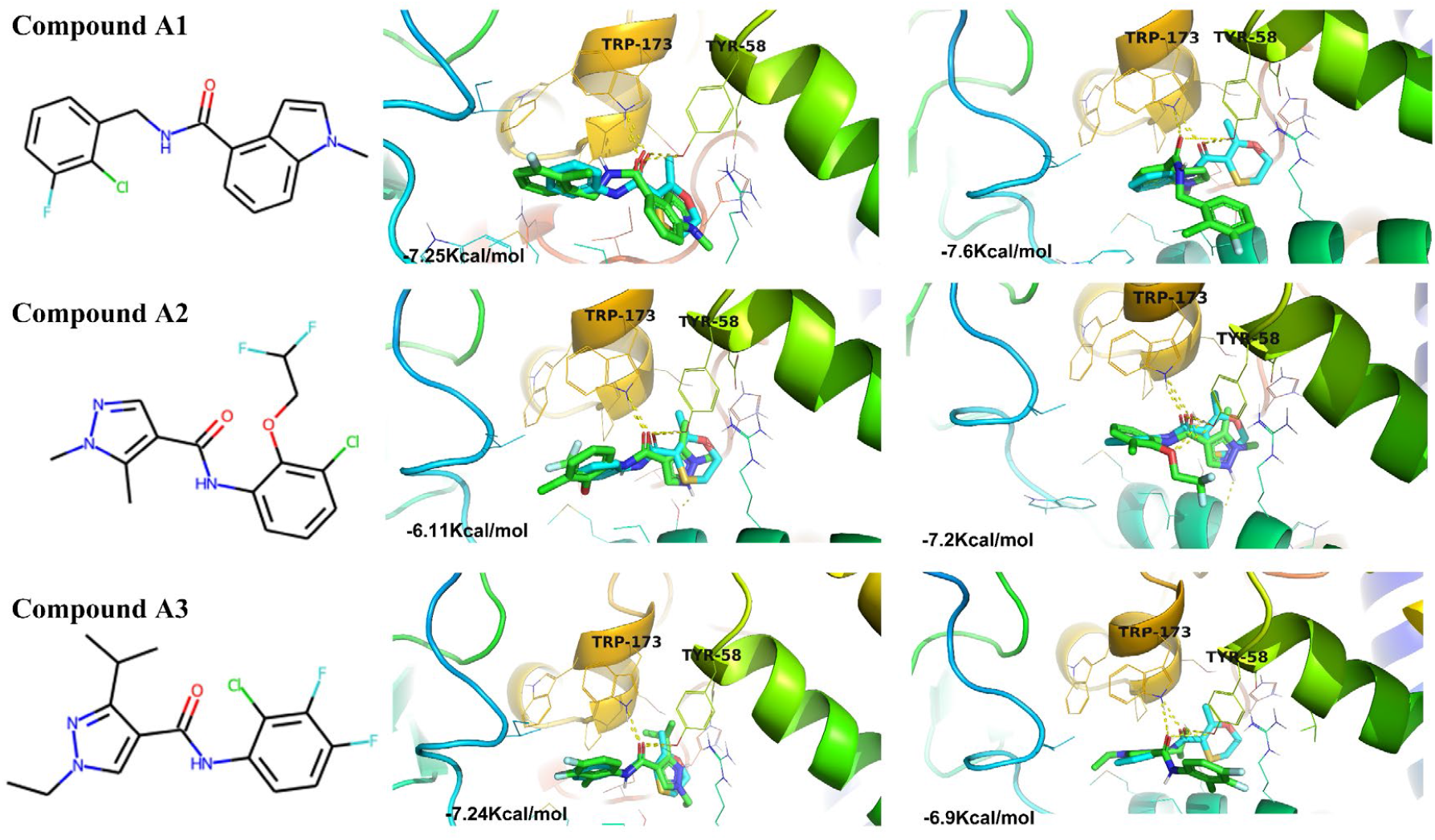
Molecular docking results. Figures show the structures and docking conformations of three molecules **A1-A3** selected from the generated molecules for subsequent synthesis and bioactivity evaluation. The left part is the structure of **A1-A3**, the middle part is the autodock docking result, and the right partis the vina docking result.

The three selected molecules **A1-A3** (Fig. 3) from 91961 generated molecules through a series of screenings for subsequent antifungal activity evaluation. The generated compound which has apparent inhibitory effect was subsequently subjected to two rounds of optimization. The compounds **A1-A3** and all subsequent structurally optimized compounds were synthesized by using a similar synthesis route and method as shown in Fig. 4 [36-37]. The specific synthesis steps and spectral data characterizing the structures of each target compound have been listed in the Supplementary Information.

**Fig. 4.**
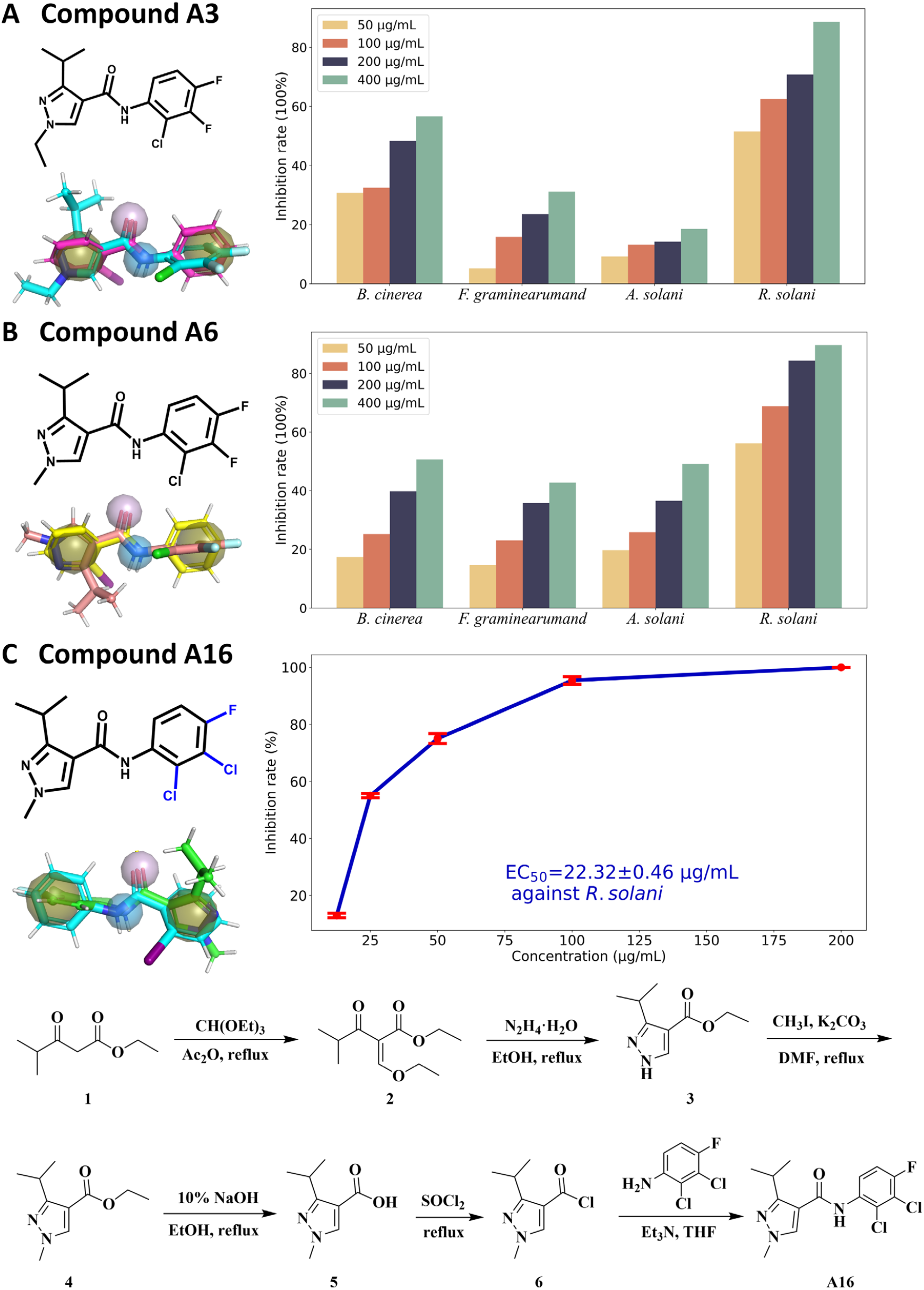
Structures and antifungal activity information of compounds A3, A6 and A16. The analyses of structure, pharmacophore, fungal inhibitory ability for the best compound **A3** initially selected for synthesis (A), the best compound **A6** found after the first round of optimization (B), and the best compound **A16** found after the second round of optimization (C). The brown sphere in the pharmacophore picture is the hydrophobic center, the blue sphere is the hydrogen bond donor, and the purple sphere is the hydrogen bond acceptor. The purple molecule in figure A, the yellow molecule in figure B, and the blue molecule in figure C are all benodanil, and the other molecules are generated molecule or the optimized molecule.

### Antifungal activity of target compounds A1-A21 and structure activity relationship

The inhibitory effects of the target compounds **A1-A3** on *Rhizoctonia solani* (*R. solani*), *Botrytis cinerea* (*B. cinerea*), *Fusarium graminearum* (*F. graminearum*) and *Alternaria solani* (*A. solani*) at a concentration of 100 μg/mL were first tested using the mycelial growth rate method with benodanil as the positive control. The antifungal activity of compounds **A1-A3** is shown in Table 1. The compounds **A1** and **A2** showed week inhibitory effect on *R. solani, B. cinerea* and *F. graminearum*and. Encouragingly, the compound **A3** was very prominent, its inhibitory rate against *R. solani* reached 62.50%. Its antifungal effect at other concentrations is shown in Fig. 4A. The pharmacophore analysis of the **A3** molecule was conducted, and it was found that the **A3** molecule, like the benodanil molecule, has two hydrophobic centers, a hydrogen bond acceptor and a hydrogen bond donor.

**Table 1.**
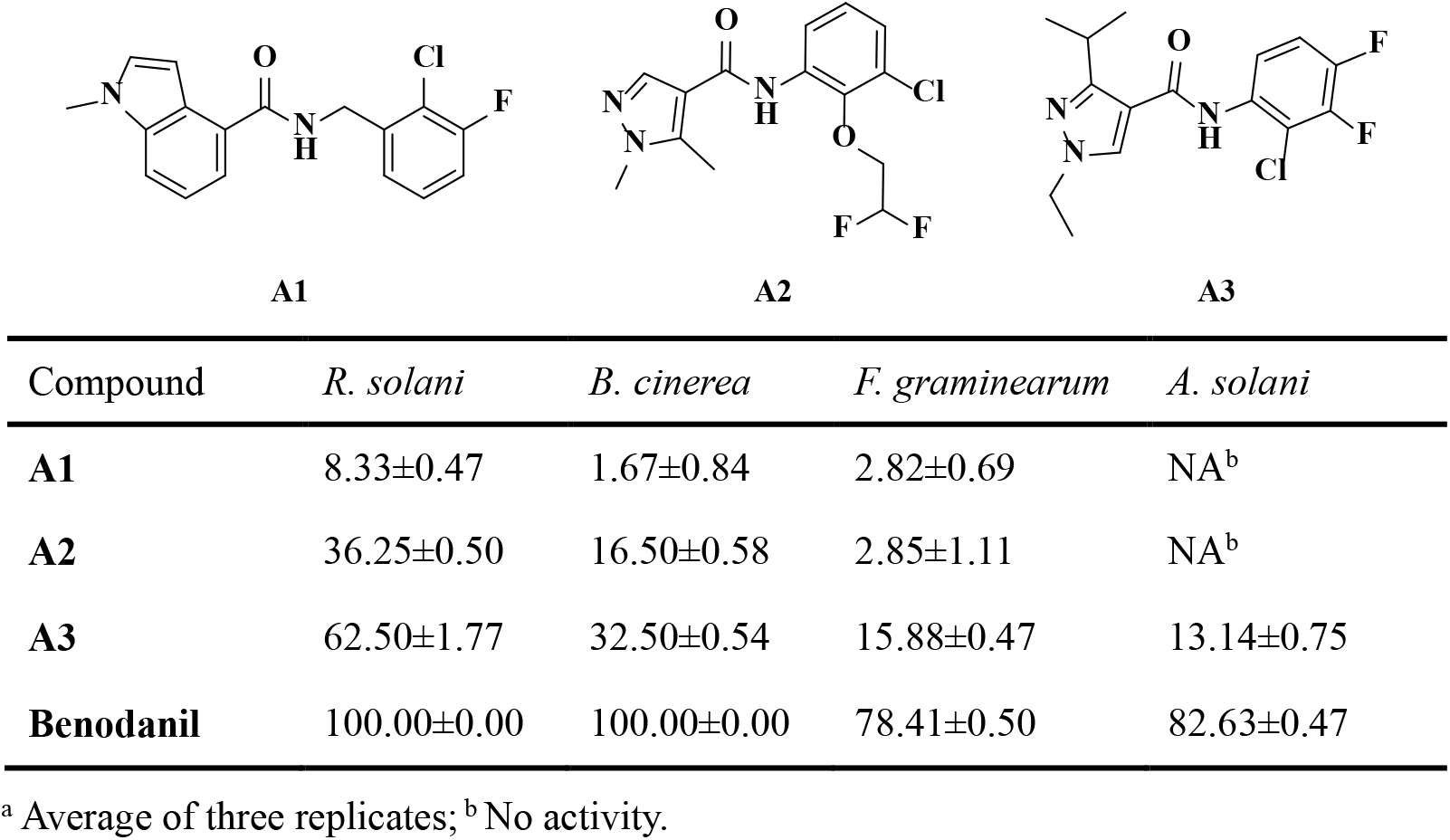
Antifungal activity of initially selected compounds A1-A3 at 100 μg/mL^a^.

Based on the result in Table 1, the substituents at 1-position and 3-position of the pyrazole ring were first optimized with **A3** as the lead compound. As can be seen from Table 2, when the isopropyl at 3-position of pyrazole ring was replaced by ethyl or cyclopropyl, the obtained compounds **A4** and **A5** showed a decrease in inhibition against *R. solani, B. cinerea* and *F. graminearum*and at the concentration of 100 μg/mL, and a pattern of **A3** > **A4** > **A5** appeared. When the ethyl at 1-position of pyrazole ring was replaced by methyl or isopropyl, the obtained compounds **A6** and **A7** respectively showed an increasing and then decreasing inhibitory effect on *R. solani* at the concentration of 100 μg/mL. The pharmacophore analysis of the **A6** molecule was conducted, and it was also found that the **A6** molecule, like the benodanil molecule, has two hydrophobic centers, a hydrogen bond acceptor and a hydrogen bond donor. Overall, when the pyrazole ring contains a methyl at the 1-position and an isopropyl at the 3-position, which plays a key role in improving the activity of the target compounds.

**Table 2.** Antifungal activity of compounds A4-A7 optimized in first round at 100 μg/mL^a^.

| Compound | *R. solani* | *B. cinerea* | *F. graminearum* | *A. solani* |
| --- | --- | --- | --- | --- |
| **A4** | 35.54±0.19 | 15.40±0.69 | 9.62±0.49 | NA<sup>b</sup> |
| **A5** | 23.33±0.47 | 8.81±1.70 | 6.12±0.37 | 2.72±0.50 |
| **A6** | 68.83±0.82 | 25.18±0.74 | 23.05±0.58 | 25.87±0.76 |
| **A7** | 47.52±1.24 | 22.42±0.50 | 12.91±0.47 | 13.68±0.58 |
| **Benodanil** | 100.00±0.00 | 100.00±0.00 | 79.72±1.12 | 82.37±0.37 |

<sup>a</sup> Average of three replicates; <sup>b</sup> No activity.

### Antifungal activity of optimized compounds A8-A21 in second round

Based on the result in Table 2, the substituents at the benzene ring were further optimized using **A6** as the lead compound. As can be seen from Fig.5, Table 3 and Table 4, the inhibitory effect of compounds **A8**-**A21** on *R. solani* was significantly better than their inhibitory effect on *B. cinerea, F. graminearum*and and *A. solani*. The target compounds **A8** (R=2-Cl-4-F-6-Me), **A11** (R=3-*i*-PrO-4-F) and **A12** (R=2-Cl-4-F-5-Me) were successfully synthesized to further explore the inhibitory effect upon the introduction of alkyl and alkoxy groups into the benzene ring. Among them, the compounds **A8** and **A11** showed the most significant inhibitory effect against *R. solani* with 73.25% and 80.57% inhibition, respectively. Fixing the group at 2-position of the benzene ring to be a Cl atom and the group at 4-position to be a F atom, and making changes to the other positions, the compounds **A8, A14** (R=2-Cl-4-F), **A12, A16** (R=2,3-Cl_2_-4-F) and **A21** (R=2-Cl-4,5-F_2_) were obtained. The compounds **A16** and **A21** inhibited *R. solani* fungus by more than 80%, which was significantly better than **A8** (73.25%), **A12** (37.27%) and **A14** (78.23%). Among them, the compounds **A16** and **A21** achieved EC_50_ values of 22.32 μg/mL and 42.02 μg/mL against *R. solani*, respectively. For fixed benzene rings with F atoms at positions 3 and 4, changes to other positions gave the compounds **A13** (R=2-Br-3,4-F_2_) and **A17** (R=3,4-F_2_). The compound **A13** (EC_50_= 49.41 μg/mL) showed better inhibitory effect against *R. solani* than compound **A17**. For fixed benzene rings with F atoms at positions 2 and 4, changes to other positions gave the compounds **A9** (R=2,4-F_2_-3-Cl) and **A15** (R=2,4-F_2_). The compound **A9** (EC_50_= 39.41 μg/mL) showed better inhibitory effect against *R. solani* than compound **A15**. The compounds **A10** (R=2,3-F_2_-4-Cl), **A18** (R=2-F-3,4-Cl_2_), **A19** (R=2,3,4,5-F_4_) and **A20** (R=4-Cl-3,5-F_2_) were obtained by changing the substituents at other positions of the benzene ring. The compounds **A18** and **A19** exhibited more than 80% inhibition on *R. solani*, which was significantly better than **A10** (73.25%) and **A20** (37.27%). Among them, the compounds **A18** and **A19** achieved EC_50_ values of 26.56 μg/mL and 49.01 μg/mL against *R. solani*, respectively. Above all, this second round of structural optimization resulted in the compound **A16**, which showed remarkable inhibitory activity against phytopathogenic fungi. As potential succinate dehydrogenase inhibitors, **A**-series compounds provide a unique molecular scaffold for the development of new fungicides.

**Fig. 5.**
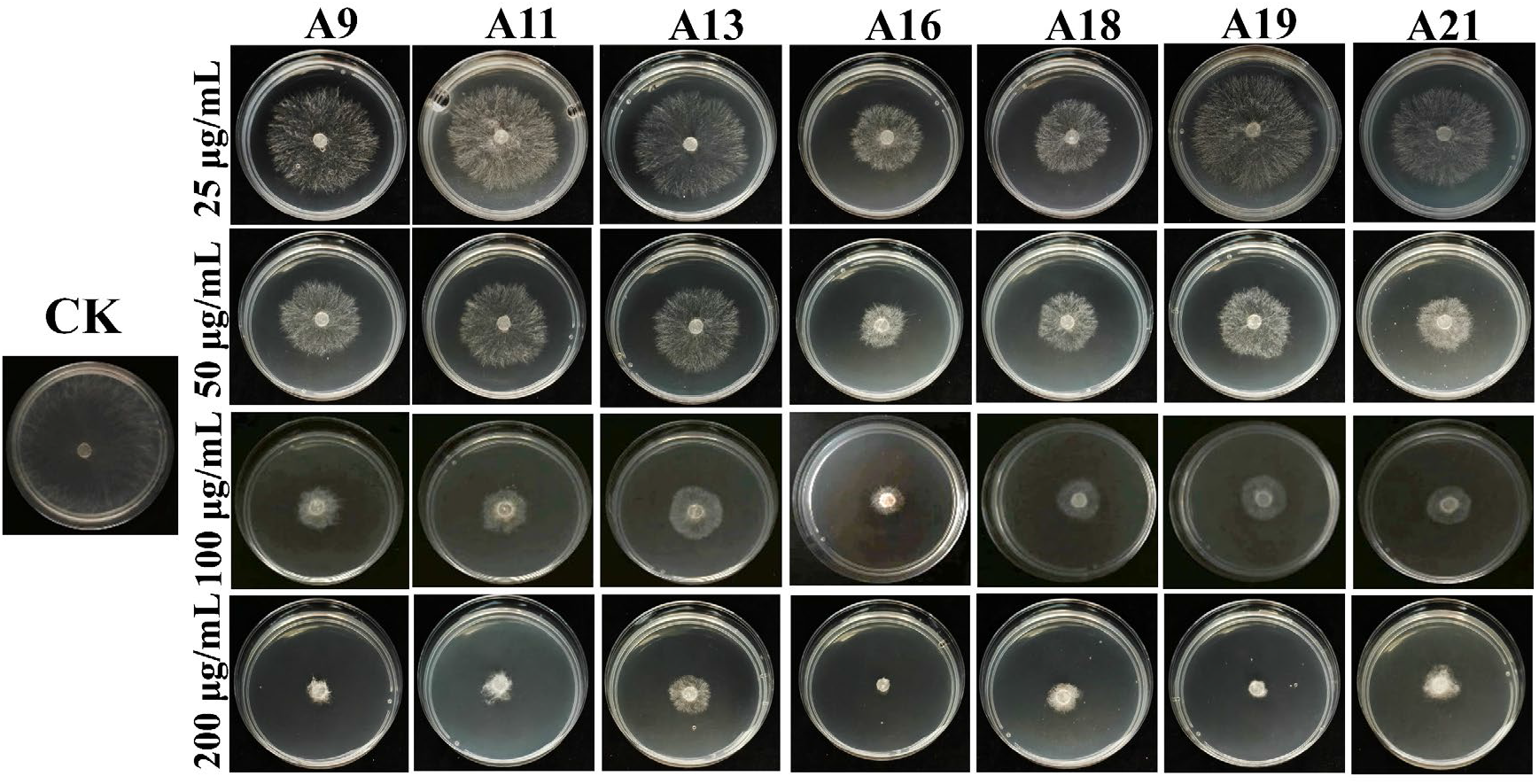
Inhibition effect diagram of some target compounds on mycelium of *R. solani*.

**Table 3.** Antifungal activity of compounds A8-A21 optimized in second round at 100 μg/mL^a^.

| Compound | R | *R. solani* | *B. cinerea* | *F. graminearum* | *A. solani* |
| --- | --- | --- | --- | --- | --- |
| **A8** | 2-Cl-4-F-6-Me | 73.25±0.19 | 49.12±1.38 | 2.38±0.63 | 40.54±0.18 |
| **A9** | 2,4-F<sub>2</sub>-3-Cl | 87.82±1.43 | 80.15±2.21 | 54.07±1.21 | 63.36±0.26 |
| **A10** | 2,3-F<sub>2</sub>-4-Cl | 71.22±0.20 | 48.49±0.69 | 36.34±1.60 | 69.79±1.11 |
| **A11** | 3-*i*-PrO-4-F | 80.57±0.27 | 81.78±0.14 | 26.31±1.17 | 27.73±0.78 |
| **A12** | 2-Cl-4-F-5-Me | 37.27±0.18 | 34.04±1.19 | 63.89±0.57 | 32.67±0.34 |
| **A13** | 2-Br-3,4-F<sub>2</sub> | 81.32±0.87 | 76.13±0.09 | 65.23±1.29 | 57.09±0.73 |
| **A14** | 2-Cl-4-F | 78.23±0.81 | 81.71±1.18 | 80.78±1.08 | 59.40±0.73 |
| **A15** | 2,4-F<sub>2</sub> | 51.10±0.54 | 29.02±0.79 | 17.85±1.06 | 52.39±0.66 |
| **A16** | 2,3-Cl<sub>2</sub>-4-F | 95.39±0.12 | 90.89±0.28 | 75.77±0.66 | 82.17±0.86 |
| **A17** | 3,4-F<sub>2</sub> | 70.96±0.69 | 32.16±0.59 | 15.43±1.18 | 55.76±0.23 |
| **A18** | 2-F-3,4-Cl<sub>2</sub> | 90.54±0.09 | 60.42±0.95 | 58.40±0.75 | 76.24±1.45 |
| **A19** | 2,3,4,5-F<sub>4</sub> | 84.96±0.09 | 58.54±0.55 | 82.15±0.27 | 63.34±0.94 |
| **A20** | 4-Cl-3,5-F<sub>2</sub> | 76.85±0.42 | 57.27±0.26 | 43.62±1.93 | 64.35±0.48 |
| **A21** | 2-Cl-4,5-F<sub>2</sub> | 88.89±0.38 | 83.35±0.19 | 76.46±0.50 | 58.74±1.23 |
| **Benodanil** | - | 100.00±0.00 | 100.00±0.00 | 77.72±0.58 | 81.68±0.47 |

<sup>a</sup> Average of three replicates.

**Table 4.** Antifungal EC_50_ values of some target compounds against *R. solani* ^a^.

| Compound | Regression equation | r | EC<sub>50</sub> (µg/mL) |
| --- | --- | --- | --- |
| **A9** | $y = 2.28x + 1.35$ | 0.9806 | $39.41 \pm 0.03$ |
| **A11** | $y = 1.88x + 1.81$ | 0.9605 | $49.68 \pm 0.68$ |
| **A13** | $y = 2.19x + 1.28$ | 0.9855 | $49.41 \pm 1.12$ |
| **A16** | $y = 2.43x + 1.72$ | 0.9958 | $22.32 \pm 0.46$ |
| **A18** | $y = 2.31x + 1.71$ | 0.9943 | $26.56 \pm 0.93$ |
| **A19** | $y = 2.51x + 0.75$ | 0.9834 | $49.01 \pm 0.67$ |
| **A21** | $y = 2.46x + 0.99$ | 0.9724 | $42.02 \pm 1.54$ |
| **Benodanil** | $y = 0.93x + 4.26$ | 0.9990 | $6.24 \pm 0.48$ |

<sup>a</sup> Average of three replicates.

### Effect of Compound A16 on Mycelial Morphology

The effect of target compound **A16** on mycelial growth and morphology of *R. solani* was investigated by scanning electron microscope (SEM). As shown in Fig. 6, the mycelial morphology of the blank control group is uniform, smooth, regular and complete. After treatment with **A16** at 100 μg/mL, the mycelia were observed being universally wrinkled and collapsed. These observed phenomena indicated that compound **A16** caused serious damage to the mycelial cell membrane and inhibited the growth and development of mycelia of *R. solani*.

**Fig. 6.**
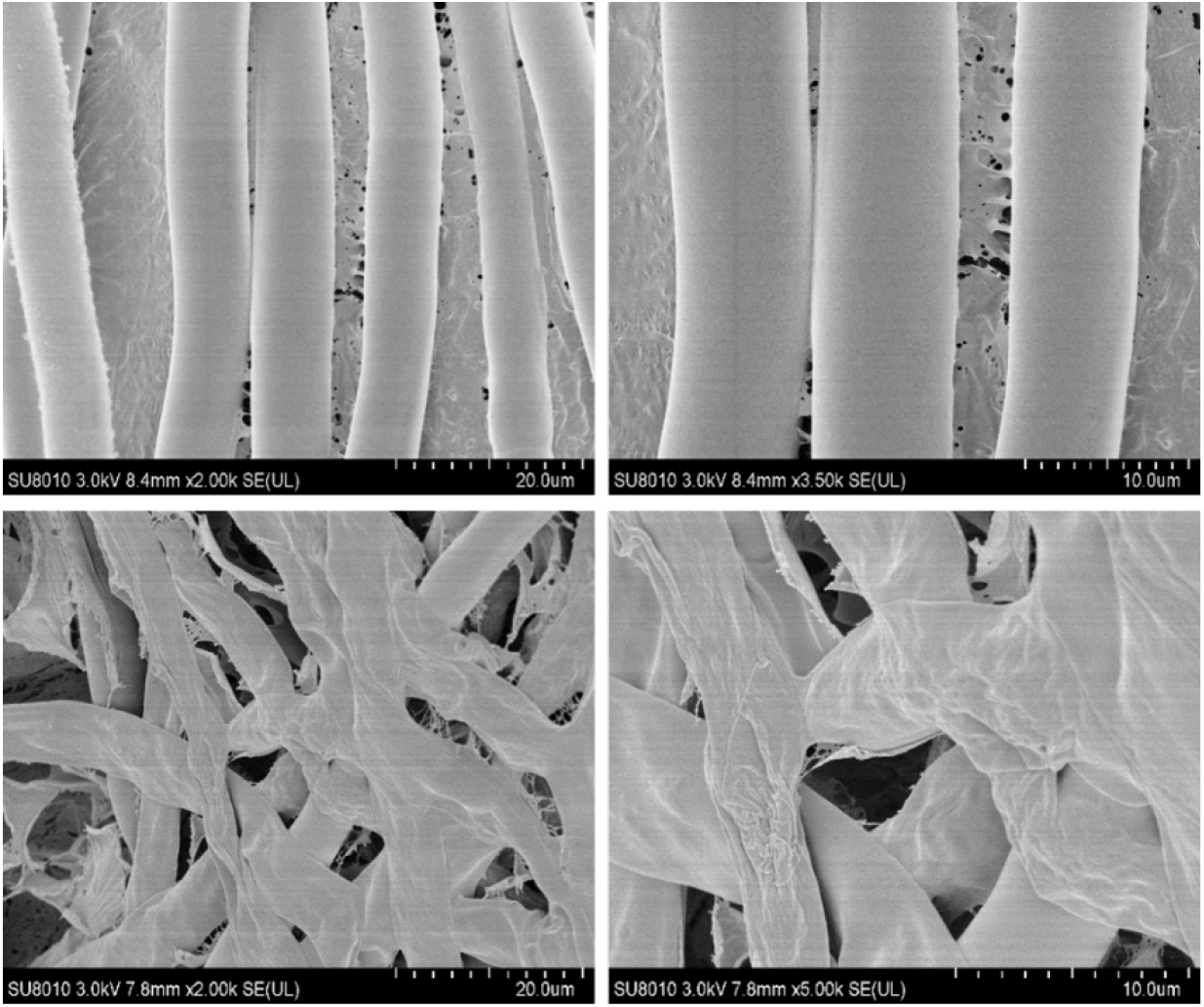
Effect of compound A16 on mycelial morphology.

### Inhibitory Effect against Fungal SDH *in vivo*

In order to verify whether SDH is the potential target enzyme for the target compounds, the compounds **A16** and **A18** with excellent antifungal effect were selected to determine the inhibitory activity against SDH *in vivo* of *R. solani*. Benodanil was used as the positive controls. The results of enzyme activity test (Table 5) showed that **A16** had obvious inhibitory activity against succinate dehydrogenase of *R. solani*, and its IC_50_ value reached 110.39 μg/mL, which was relatively close to that of benodanil (72.60 μg/mL).

**Table 5.**
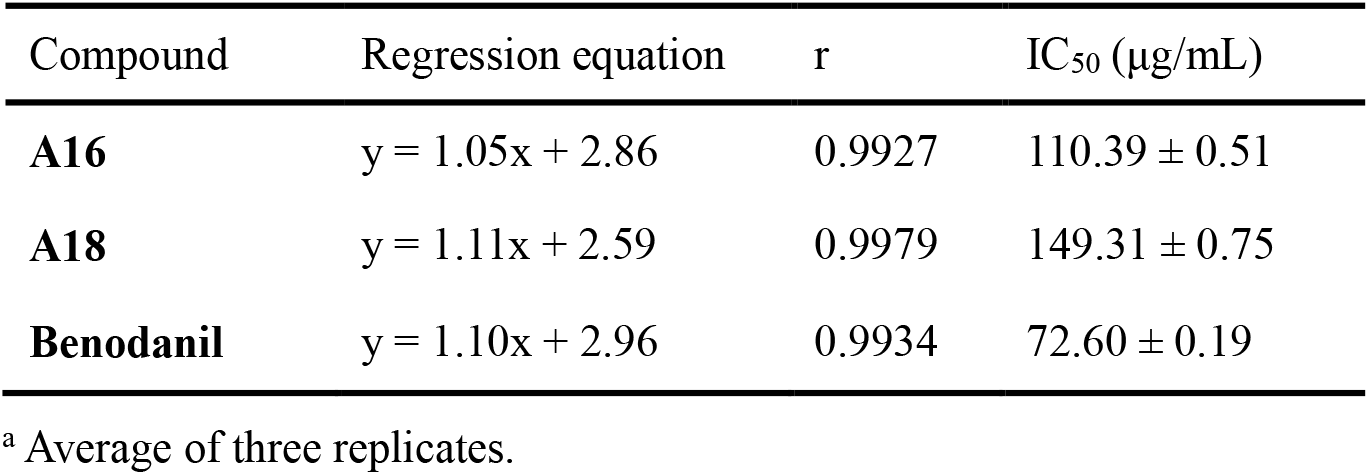
IC_50_ values of some target compounds against SDH from *R. solani* ^a^.

## Discussion

New fungicide development is an intricate endeavor, demanding expertise across diverse disciplines and knowledge domains. Consequently, fungicide research and development are characterized by high investment, lengthy development cycles, and significant risk. Statistics indicate that launching a new fungicide variety typically requires $300 million and 12 years, with a concerning trend of increasing costs.

Identifying promising compounds with desired biological activity represents the initial step in this process. The most common approach for discovering novel candidates involves high-throughput screening of existing compound databases. While hit compounds often possess unique chemical structures and promising activities, the inherent limitations in diversity within existing databases can lead to suboptimal screening outcomes. This article addresses this challenge by employing a trained Transformer model to generate a customized virtual compound library for succinate dehydrogenase inhibitors. The generated molecules exhibit high validity, novelty, and structural uniqueness, paving the way for a diverse and promising library. Subsequent screening of this library yielded three readily synthesizable compounds for further evaluation. Following two rounds of structural optimization focused on the pyrazole and benzene rings, the most active candidate ultimately yielded potential succinate dehydrogenase inhibitors with demonstrable inhibitory effects.

Traditional approaches to designing succinate dehydrogenase inhibitors rely heavily on experience-driven modification of existing structures or database-driven virtual screening. This work represents the first attempt to leverage generative models in this domain, opening up a new avenue for the design and development of novel succinate dehydrogenase inhibitors and potentially fungicides in general. While the activity of our screened compounds falls short of the existing benchmark, benodanil, it highlights the potential for further improvement. Firstly, the model itself can be enhanced by integrating activity prediction methods or incorporating structural information of pathogenic fungal succinate dehydrogenase predicted through homology modeling or AlphaFold. This would enable the structure-based drug design. Secondly, addressing the high resistance risk associated with existing inhibitors is crucial. Designing molecules with low cross-resistance or targeting new binding sites presents an avenue to overcome this challenge. Moving forward, we will focus on exploring these two aspects, utilizing both predicted structures of pathogenic fungi and scaffold hopping strategies to inform future molecular design efforts.

In conclusion, our study showcases the potential of generative models in the development of novel succinate dehydrogenase inhibitors, providing a new thought for a more efficient and diverse approach to fungicide discovery.

## Methods

### Data preparation

MOSES is a benchmark platform for molecular generative models, and the data used by the MOSES benchmark platform was used to pretrain the model [38]. The dataset used in MOSES is over 1.9 million molecules from the Zinc dataset [39], with molecular weights ranging from 250 to 350 Da, the number of rotatable bonds below 7, and XlogP below 3.5. The data of succinate dehydrogenase inhibitors came from the C2 group in the document provided by the fungicide resistance action committee in 2020, with a total of 23 molecules. The SMILES data of the molecules were searched in the PubChem database [40]. SMILES data processing was as follows: use python’s OpenBabel package to read the molecular data in SMILES format [41], remove hydrogen, extract the atom type sequence, obtain the index of atoms which connect the bond and bond type. The atomic index and bond type are used to construct the adjacency matrix.

### Implementation of generated model

The model used in this article is Transformer, as well as its input is the atom type sequence and adjacency matrix obtained in the data preparation part. The model is mainly composed of an encoder with a multi-head self-attention mechanism, an atomic decoder (FC_ATOM_) and an edge decoder (FC_BOND_). FC_ATOM_ responsible for predicting the atom type. FC_BOND_ responsible for combining the predicted atoms with the atomic sequence generated by the model to jointly predict the next column of the adjacency matrix. However, the process is somewhat different in training and generation. During the training process, this is done by encoding the corresponding atoms in the real sequence and stacking them with the existing encoding sequence to input FC_BOND_ which predicted bond type. During the generation process, the predicted bond type of FC_BOND_ is entered by encoding the predicted atoms and stacking them with existing encoding sequences. We first pre-trained the model using the data used by the MOSES benchmark platform. Secondly, the model was fine-tuned using the collected 23 succinate dehydrogenase inhibitors. This process was implemented by updating the parameters using the collected 23 succinate dehydrogenase inhibitors, based on the parameters of the converged pre-trained model using the data used by the MOSES benchmark platform. Finally, the trained model was used to generate 100,000 molecules in an autoregressive manner. The loss of this model consists of two parts, one is the cross-entropy between the predicted atom and the corresponding next atom in the sequence of atoms, and the other is the negative log-likelihood of the bond probability distribution. The parameters of the generative model were then updated using adam optimizer with learning rate of 0.0001 until convergence.

### Screening process

We used the generative model to generate 100,000 molecules and performed the following screening process. First, after removing some invalid compounds (cannot be converted from adjacency matrix to molecules, salt compounds), a total of 92,567 compounds were found. Second, after removing duplicate compounds, a total of 91,961 compounds were found. Third, after selecting molecules with amide bond structures, a total of 72,319 molecules were found. The reason for this step of screening is that the amide bond is part of the chemical structure of the classic SDHI fungicide [42]. Fourth, after selecting compounds with F atoms and Cl atoms, a total of 1,188 compounds were found. The reason for this step of screening is that SDHIs recently added by the FRAC all have F or Cl atoms. Finally, molecules with a crotonamide scaffold were selected, totaling 77 compounds. After the results and experience of molecular docking, three molecules were finally selected for subsequent evaluation of antifungal activity.

### Metrics

Three indicators (validity, novelty and uniqueness) commonly used in molecule generation tasks are used to evaluate the ability of model to learn and generate molecules [43].

Validity: The ratio of generated valid molecules to the total number of generated molecules. Invalid molecules refer to molecules that cannot be converted from the adjacency matrix and salt compounds. The calculation is as follows:

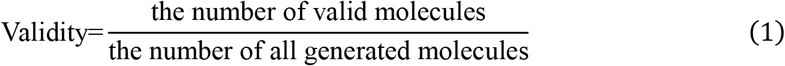

Uniqueness: The proportion of unique molecules among the generated valid molecules. If the generated molecule is unique, it means that there are no other molecules with the same structure. Calculated as follows:

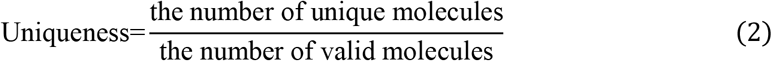

Novelty: the proportion of generated valid molecules that were not present in the original data set among the total generated valid molecules. Low novelty is a sign of overfitting. Calculated as follows:

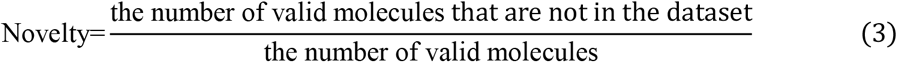

### Similarity calculation

The molecular fingerprint of the molecule was calculated using the RDKit package of Python. The molecular fingerprint is a binary vector of length 2048. The evaluation method used to calculate the similarity of two compounds is the tanimoto coefficient. The tanimoto coefficient between two molecules A and B can be calculated using molecular fingerprints through the following formula:

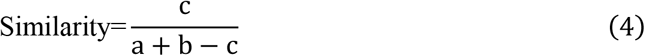

Among them, a is the number of 1 in A molecule, b is the number of 1 in B molecule, and c is the number of 1 both in A molecule and B molecule.

### Molecular properties

This article mainly evaluates the following 6 properties of the generated molecules, and uses the RDKit package in python to calculate.

Molecular weight: molecular weight is the sum of the relative atomic masses of the individual atoms in a chemical formula, usually measured in Daltons (Da).

Logp [44]: the partition coefficient compares the solubility of a solute in an equilibrium solvent. If one solvent is water and the other is oil, logp is the oil-water partition coefficient, which reflects the hydrophilicity and hydrophobicity of the chemical molecule.

Quantitative estimate of drug-likeness (QED) [45]: the QED is a method of quantifying drug similarity as a numerical value between 0 and 1.

Synthetic accessibility (SA) [46]: the SA used to measure the ease of molecule synthesis. It evaluates the ease of molecule synthesis with a numerical value between 1 and 10. The closer to 1, the easier it is to synthesize, and the closer to 10, the more difficult it is to synthesize.

Natural product-likeness (NP) [47]: the NP measures the similarity of compounds and natural products in chemical space. The NP similarity score ranges from -5 to 5, with higher numbers being more likely to be natural products.

Relative topological surface area (TPSA) [48]: the TPSA is one of the indicators describing the chemical properties of molecules. It is used to describe properties such as solubility and permeability of molecules. TPSA is obtained by calculating the surface area of a molecule, which is the area of interaction between the molecule and the solvent on the surface of the molecule.

### Molecular docking

The crystal structure of chicken heart SDH (PDB: 2FBW) [49] was selected as the receptor. The crystal structure was downloaded from the PDB database [50], and the AutoDock program [51] and Vina program [52] were used for molecular docking. The grid center is set according to the commercial succinate dehydrogenase inhibitor in 2FBW (centre_x:14.972, center_y:16.444, center_z:8.694), the AutoDock grid size is set to 40×40×40, and the Vina grid size is set to 15×15×15. The docking number of AutoDock is 20, and the docking number of Vina is 9.

### Pharmacophore analysis

Pharao [53], also known as align-it, was used for pharmacophore analysis. The pharmacophores of SDHIs were extracted separately. Using the SDHIs as reference molecules, the pharmacophore shared by SDHIs and generated molecules were extracted.

### Synthesis of target compounds

The synthesis methods of the target compounds are listed in detail in the Supplementary Information. The compound A16 was took as a representative to describe its synthesis method as follows. As shown in Fig. 4C, ethyl 4-methyl-3-oxopentanoate **1** (100.00 mmol) and triethyl orthoformate (200.00 mmol) were added to acetic anhydride (45 mL). The mixed solution was heated to reflux and stirred for 7 h. The acetic anhydride was then removed by distillation under reduced pressure to obtain intermediate **2**. Which was added to the ethanol solution (40 mL) of hydrazine hydrate (300.00 mmol) under ice bath condition and stirred for 30 min. The reaction mixture was refluxed for 5 h, and the excess ethanol was then removed by distillation under reduced pressure. The obtained residue was added to ice water (100 mL), filtered and dried under vacuum to give the intermediate **3**. Intermediate **3** (50.00 mmol), potassium carbonate (50.00 mmol) and iodomethane were added to DMF (25 mL) sequentially. At the end of the reaction, the organic phase was washed with saturated ammonium chloride solution for 5 times, the organic phase was retained, dried over anhydrous sodium sulfate, filtered, and the excess solvent was removed under reduced pressure to give intermediate **4**. Intermediate **4** (30.00 mmol) and 10% sodium hydroxide solution were added to ethanol (45 mL), heated to reflux and reacted for 2 h. The reaction solution was concentrated under vacuum to remove excess ethanol, poured into ice water, adjusted the acidity with concentrated hydrochloric acid to pH=2.0, filtered and dried to obtain the intermediate **5**. The compound **5** (5.00 mmol) was added to dichlorosulfoxide (15 mL) and heated to reflux. The reaction was carried out for 4 h. The excess dichlorosulfoxide was removed by distillation under reduced pressure to obtain the intermediate **6**. The intermediate **6** (5.00 mmol), Et_3_N (15.00 mmol) and 2,3-dichloro-4-fluoroaniline (5.00 mmol) were added sequentially to THF (15 mL) under ice bath condition. After stirring for **7** h at room temperature, the solvent was removed under reduced pressure. The obtained crude product was purified using silica gel column chromatography with petroleum ether/ethyl acetate (V/V, 2:1) as the eluent to give the target compound **A16**.

### Determination of antifungal activity *in vitro*

The mycelial growth rate method was applied to test the inhibitory effects of the target molecules against *Rhizoctonia solani* (*R. solani*), *Botrytis cinerea* (*B. cinerea*), *Fusarium graminearum* (*F. graminearum*) and *Alternaria sonali* (*A. sonali*) [54-57], which were supplied by Jiangsu Key Laboratory of Pesticide Science, Nanjing Agricultural University. Each tested compound was solubilized in same volume of DMSO and mixed with the molten Potato Dextrose Agar (PDA) to obtain the medicated medium at 100 μg/mL with three replicates. The fungus mycelia cakes (5 mm) were placed in the center of PDA medium plates with a germfree inoculation needle, and then were incubated for 3-5 days at 25 °C. The SDHI fungicide benodanil was selected as the positive control. The treatments containing equal volumes of solvent DMSO were used as the blank controls. When colony sizes of blank controls reached two-thirds of the diameters of the PDA medium, their diameters were measured using the cross method. The inhibitory effects on these four fungi were calculated as follows:

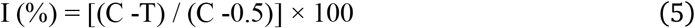

The I represents the rate of inhibition, C indicates the fungal colony diameter in the blank control, and T represents the fungal colony diameter in the chemical treatment. The standard deviation (SD) values were calculated from three repeats of inhibitory data. Other suitable concentrations of each target compound were prepared using a similar method described above. The EC_50_ values and corresponding 95% confidence intervals were calculated with DPS software 9.01.

### Determination of SDH inhibitory activity

The activated *R. solani* colonies (5 mm) were added to PDB medium (200 mL) and incubated at 25 °C for 5 days with shaking at 150 rpm. The tested compounds **A16** and **A18** and the positive control benodanil dissolved in 0.4 mL DMSO and 1.0 mL of Tween 80 were respectively added to the above PDB medium to make their final concentrations of 400.00 μg/mL, 200.00 μg/mL, 100.00 μg/mL, 50 μg/mL, 25 μg/mL, respectively, and incubated for 24 h. The treatments with equal amount of DMSO and Tween 80 were used as the blank controls. Each sample was repeated three times. Subsequent steps were performed according to the previously reported methods and the protocol on the SDH assay kit (Solebro, BC0955) [58-60].

### Microscopic Morphology Observation of *R. solani* Hypha

*R. solani* colonies with length of 4 mm and width of 4 mm were obtained from PDA medium containing 100 μg/mL of **A16**. The treatment with equal volume of the solvent DMSO (0.1 mL) was used as the blank control. The images were obtained using SU8010 scanning electron microscope (Hitachi, Japan) [61-63].

## Data availability

Source data of this article are found at https://github.com/xueyuanyuan0410/SDHIs_generation/tree/main/data

## Code availability

Source codes of this article are found at https://github.com/xueyuanyuan0410/SDHIs_generation/tree/main

## Acknowledgements

We thank Jiangsu Key Laboratory of Pesticide Science, Nanjing Agricultural University for help.

## Author Contributions

C.P., C.L.Y., Y.Z. and J.Q.C. conceived and designed the study; Y.Z., J.Q.C., L.L. and W.Q.Z. performed the experiments, analyzed the data, prepared figures and tables; C.P., C.L.Y., Y.Z., J.Q.C., Y.Y.C. and L.Y.Z. authored or reviewed drafts of the paper, and approved the final draft.

## Funding

This work was supported in part by the National Natural Science Foundation of China [ZX2200521].

## Conflict of interest

The authors declare that they have no competing interests.

## Notes

### Competing Interest Statement

The authors have declared no competing interest.

## References

[1] Huang, Y. H. et al. Discovery of N-methoxy-(biphenyl-ethyl)-pyrazole-carboxamides as novel succinate dehydrogenase inhibitors. J. Agr. Food Chem. 70, 14480–14487 (2022).

[2] Yin, Y. M., Sun, Z. Y., Wang, D. W. & Xi, Z. Discovery of benzothiazolylpyrazole-4-carboxamides as potent succinate dehydrogenase inhibitors through active fragment exchange and link approach. J. Agr. Food Chem. 71, 14471–14482 (2023).

[3] Li, H., Liu, Z., Dong, Y., Wang, Y. X. & Zhu, X. L. Design, synthesis, and fungicidal evaluation of novel N-methoxy pyrazole-4-carboxamides as potent succinate dehydrogenase inhibitors. J. Agr. Food Chem. 71, 2610–2615 (2023).

[4] Li, H., Wang, Y. X., Zhu, X. L. & Yang, G. F. Discovery of a fungicide candidate targeting succinate dehydrogenase via computational substitution optimization. J. Agr. Food Chem. 69, 13227–13234 (2021).

[5] Wang, W. et al. Rational Design, synthesis, and biological evaluation of fluorine- and chlorine-substituted pyrazol-5-yl-benzamide derivatives as potential succinate dehydrogenase inhibitors. J. Agr. Food Chem. 70, 7566–7575 (2022).

[6] Jiang, W. et al. Design, synthesis, inhibitory activity, and molecular modeling of novel pyrazole-furan/thiophene carboxamide hybrids as potential fungicides targeting succinate dehydrogenase. J. Agr. Food Chem. 71, 729–738 (2023).

[7] Weininger & David. SMILES, a chemical language and information system. 1. introduction to methodology and encoding rules. J. Chem. Inf. Comput. Sci. 28, 31–36 (1988).

[8] Segler, M. H. S., Kogej, T., Tyrchan, C. & Waller, M. P. Generating focused molecule libraries for drug discovery with recurrent neural networks. ACS Cent. Sci. 4, 120–131 (2018).

[9] Blaschke, T. et al. REINVENT 2.0: an AI tool for de novo drug design. J. Chem. Inf. Model. 60, 5918–5922 (2020).

[10] Arus-Pous, J. et al. Exploring the GDB-13 chemical space using deep generative models. J. Cheminformatics. 11, 20 (2019).

[11] Kingma D.P. & Welling M.. Auto-encoding variational bayes. 1312.6114.

[12] Simonovsky, M. & Komodakis, N. GraphVAE: towards generation of small graphs using variational autoencoders. 1802.03480.

[13] Gómez-Bombarelli, R. et al. Automatic chemical design using a data-driven continuous representation of molecules. ACS Cent. Sci. 4, 268–276 (2018).

[14] Winter, R., Montanari, F., Noe, F. & Clevert, D. A. Learning continuous and data-driven molecular descriptors by translating equivalent chemical representations. Chem. Sci. 10, 1692–1701 (2019).

[15] Goodfellow I.J. et al. Generative adversarial nets. 1406.2661v1.

[16] Yu, L.T., Zhang, W.N., Wang, J. & Yu, Y. SeqGAN: sequence generative adversarial nets with policy gradient. 1609.05473.

[17] Guimaraes, G. L., Sanchez-Lengeling, B., Outeiral, C., Farias, P.L.C. & Aspuru-Guzik, A. Objective-reinforced generative adversarial networks (ORGAN) for sequence generation models. 1705.10843.

[18] Cao, N. D. & Kipf, T. MolGAN: an implicit generative model for small molecular graphs. 1805.11973.

[19] Prykhodko, O. et al. A de novo molecular generation method using latent vector based generative adversarial network. J. Cheminformatics. 11, 74 (2019).

[20] Vaswani, A. et al. Attention is all you need. 1706.03762.

[21] Cofala, T. & Kramer, O. Transformers for molecular graph generation. ESANN 2021;123–128 (2021).

[22] Cofala, T., Teusch, T. & O. Kramer. Spatial generation of molecules with transformers. IJCNN 2021;1–7(2021).

[23] Bagal, V., Aggarwal, R., Vinod, P. K. & Priyakumar, U. D. MolGPT: molecular generation using a transformer-decoder model. J. Chem. Inf. Model. 62, 2064–2076 (2022).

[24] Arus-Pous, J. et al. SMILES-based deep generative scaffold decorator for de-novo drug design. J. Cheminformatics. 12, 38 (2020).

[25] Lim, J., Hwang, S. Y., Moon, S., Kim, S. & Kim, W. Y. Scaffold-based molecular design with a graph generative model. Chem. Sci. 11, 1153–1164 (2019).

[26] Kaitoh, K. & Yamanishi, Y. Scaffold-retained structure generator to exhaustively create molecules in an arbitrary chemical space. J. Chem. Inf. Model. 62, 2212–2225 (2022).

[27] Zhang, O. et al. ResGen is a pocket-aware 3D molecular generation model based on parallel multiscale modelling. Nat. Mach. Intell. 5, 1020–1030 (2023).

[28] Zhang, O. et al. Learning on topological surface and geometric structure for 3D molecular generation. Nat. Comput. Sci. 3, 849–859 (2023).

[29] Zhu, H., Zhou, R., Cao, D., Tang, J. & Li, M. A pharmacophore-guided deep learning approach for bioactive molecular generation. Nat. Commun. 14, 6234 (2023).

[30] Godinez, W. J. et al. Design of potent antimalarials with generative chemistry. Nat. Mach. Intell. 4, 180–186 (2022).

[31] Yang, L. et al. Transformer-based generative model accelerating the development of novel BRAF inhibitors. ACS Omega. 6, 33864–33873 (2021).

[32] Li, Y. et al. Generative deep learning enables the discovery of a potent and selective RIPK1 inhibitor. Nat. Commun. 13, 6891 (2022).

[33] Moret, M. et al. Leveraging molecular structure and bioactivity with chemical language models for de novo drug design. Nat. Commun. 14, 114 (2023).

[34] Xiong, L. et al. Discovery of N-benzoxazol-5-yl-pyrazole-4-carboxamides as nanomolar SQR inhibitors. Eur. J. Med. Chem. 95, 424–34 (2015).

[35] Xiong, L. et al. Structure-based discovery of potential fungicides as succinate ubiquinone oxidoreductase inhibitors. J. Agr. Food Chem. 65, 1021–1029 (2017).

[36] Chai, J. Q. et al. Potential succinate dehydrogenase inhibitors bearing a novel pyrazole-4-sulfonohydrazide scaffold: molecular design, antifungal evaluation, and action mechanism. J. Agr. Food Chem. 71, 9266–9279 (2023).

[37] Wang, X. et al. Novel pyrazole-4-acetohydrazide derivatives potentially targeting fungal succinate dehydrogenase: design, synthesis, three-dimensional quantitative structure-activity relationship, and molecular docking. J. Agr. Food Chem. 69, 9557–9570 (2021).

[38] Polykovskiy, D. et al. Molecular sets (MOSES): a benchmarking platform for molecular generation models. Front. Pharmacol. 11, 565644 (2020).

[39] Sterling, T. & Irwin, J. J. ZINC 15--ligand discovery for everyone. J. Chem. Inf. Model. 55, 2324–37 (2015).

[40] Kim, S. et al. PubChem 2023 update. Nucleic Acids Res. 51, D1373–D1380 (2023).

[41] O’Boyle, N. M., Morley, C. & Hutchison, G. R. Pybel: a Python wrapper for the OpenBabel cheminformatics toolkit. Chem. Cent. J. 2, 5 (2008).

[42] Huang, Y. H. et al. Structure-based discovery of new succinate dehydrogenase inhibitors via scaffold hopping strategy. J. Agr. Food Chem. 71, 18292–18300 (2023).

[43] Skinnider, M. A., Stacey, R. G., Wishart, D. S. & Foster, L. J. Chemical language models enable navigation in sparsely populated chemical space. Nat. Mach. Intell. 3, 759–770 (2021).

[44] Wildman, S. A. & Crippen, G. M. Prediction of physicochemical parameters by atomic contributions. J. Chem. Inf. Comput. Sci.. 39, 868–873 (1999).

[45] Bickerton, G. R., Paolini, G. V., Besnard, J., Muresan, S. & Hopkins, A. L. Quantifying the chemical beauty of drugs. Nat. Chem. 4, 90–8 (2012).

[46] Ertl, P. & Schuffenhauer, A. Estimation of synthetic accessibility score of drug-like molecules based on molecular complexity and fragment contributions. J. Cheminformatics. 1, 8 (2009).

[47] Ertl, P., Roggo, S. & Schuffenhauer, A. Natural product-likeness score and its application for prioritization of compound libraries. J. Chem. Inf. Model. 48, 68–74 (2008).

[48] Ertl, P., Rohde, B. & Selzer, P. Fast calculation of molecular polar surface area as a sum of fragment-based contributions and its application to the prediction of drug transport properties. J. Med. Chem. 43, 3714–3717 (2000).

[49] Huang, L. S. et al. 3-nitropropionic acid is a suicide inhibitor of mitochondrial respiration that, upon oxidation by complex II, forms a covalent adduct with a catalytic base arginine in the active site of the enzyme. J. Biol. Chem. 281, 5965–72 (2006).

[50] Burley, S. K. et al. RCSB protein data bank (RCSB.org): delivery of experimentally-determined PDB structures alongside one million computed structure models of proteins from artificial intelligence/machine learning. Nucleic Acids Res. 51, D488–D508 (2023).

[51] Morris, G. M. et al. AutoDock4 and AutoDockTools4: automated docking with selective receptor flexibility. J. Comput. Chem. 30, 2785–91 (2009)

[52] Trott, O. & Olson, A. J. AutoDock Vina: improving the speed and accuracy of docking with a new scoring function, efficient optimization, and multithreading. J. Comput. Chem. 31, 455–61 (2010).

[53] Taminau, J., Thijs, G. & De Winter, H. Pharao: pharmacophore alignment and optimization. J. Mol. Graphics Modell. 27, 161–9 (2008).

[54] Sun, Y. et al. Design, synthesis, and fungicidal evaluation of novel 1,3-benzodioxole-pyrimidine derivatives as potential succinate dehydrogenase inhibitors. J. Agr. Food Chem. 70, 7360–7374 (2022).

[55] Zhang, A. et al. Discovery of N-(4-fluoro-2-(phenylamino)phenyl)-pyrazole-4-carboxamides as potential succinate dehydrogenase inhibitors. Pestic. Biochem. Physiol. 158, 175–184 (2019).

[56] Yan, Z. et al. Design, synthesis, DFT study and antifungal activity of the derivatives of pyrazolecarboxamide containing thiazole or oxazole ring. Eur. J. Med. Chem. 149, 170–181 (2018).

[57] Ye, Y. H. et al. Synthesis and antifungal activity of nicotinamide derivatives as succinate dehydrogenase inhibitors. J. Agr. Food Chem. 62, 4063–71 (2014).

[58] Yu, B. et al. Design, synthesis and biological evaluation of pyrazole-aromatic containing carboxamides as potent SDH inhibitors. Eur. J. Med. Chem. 214, 113230 (2021).

[59] Cheng, W. et al. Design, synthesis and inhibitory activity of novel 2, 3-dihydroquinolin-4(1H)-one derivatives as potential succinate dehydrogenase inhibitors. Eur. J. Med. Chem. 214, 113246 (2021).

[60] Wang, W. et al. Synthesis and biological activity of novel pyrazol-5-yl-benzamide derivatives as potential succinate dehydrogenase inhibitors. J. Agr. Food Chem. 69, 5746–5754 (2021).

[61] Tai, L. et al. Novel fungicidal phenylethanol derivatives linking a trifluoromethyl pyrazole pharmacophore: design, synthesis, crystal structure, and biology evaluation. New J. Chem. 47, 12850–12860 (2023).

[62] Sun, S.-X. et al. Design, synthesis, antifungal evaluation, and molecular docking of novel 1,2,4-triazole derivatives containing oxime ether and cyclopropyl moieties as potential sterol demethylase inhibitors. New J. Chem. 45, 18898–18907 (2021).

[63] Sun, S. et al. Novel (Z)/(E)-1,2,4-triazole derivatives containing oxime ether moiety as potential ergosterol biosynthesis inhibitors: design, preparation, antifungal evaluation, and molecular docking. Mol. Diversity. 27, 145–157 (2023).

