## supplemental file1 for "Design and optimization of novel succinate dehydrogenase inhibitors against agricultural fungi based on Transformer model"

#### Supplementary Information

### 1 Similarity between SDHIs and Generated Molecules

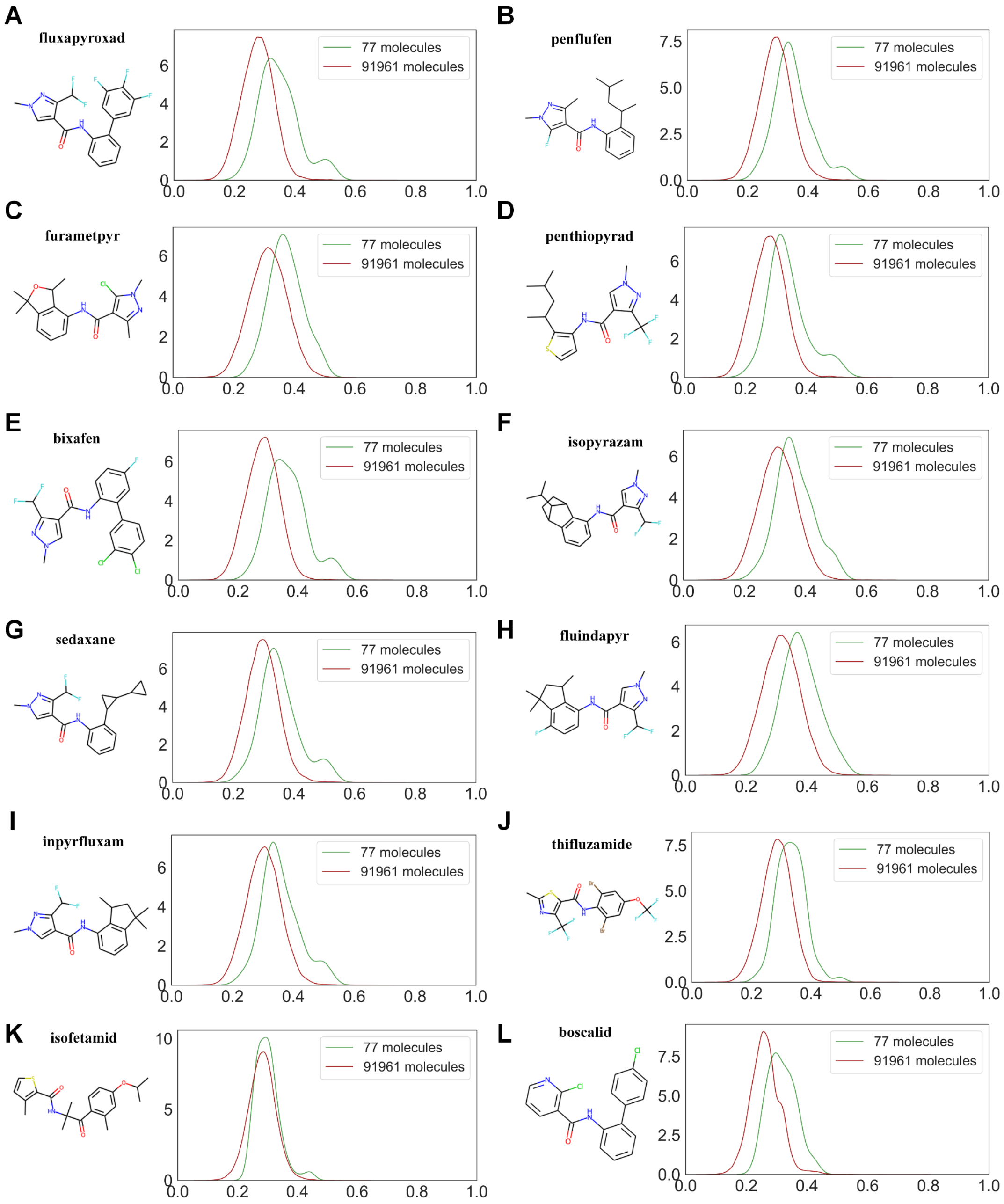


**Fig. 1| Similarity between SDHIs and generated molecules.**

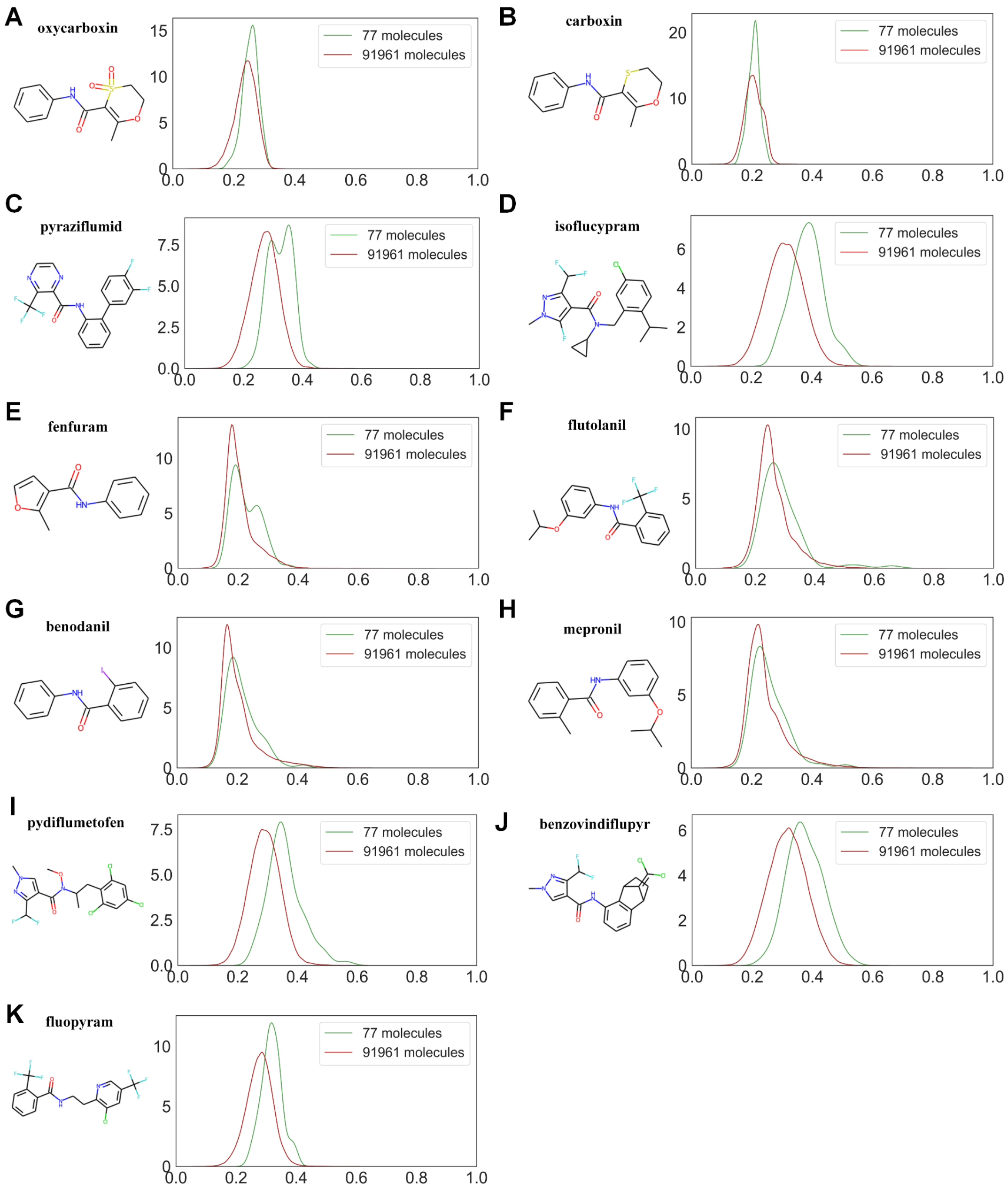


**Fig. 2| Similarity between SDHIs and generated molecules.**

#### 2 Synthesis of Target Compounds

*2.1 Synthesis of Intermediates **2a–2g***

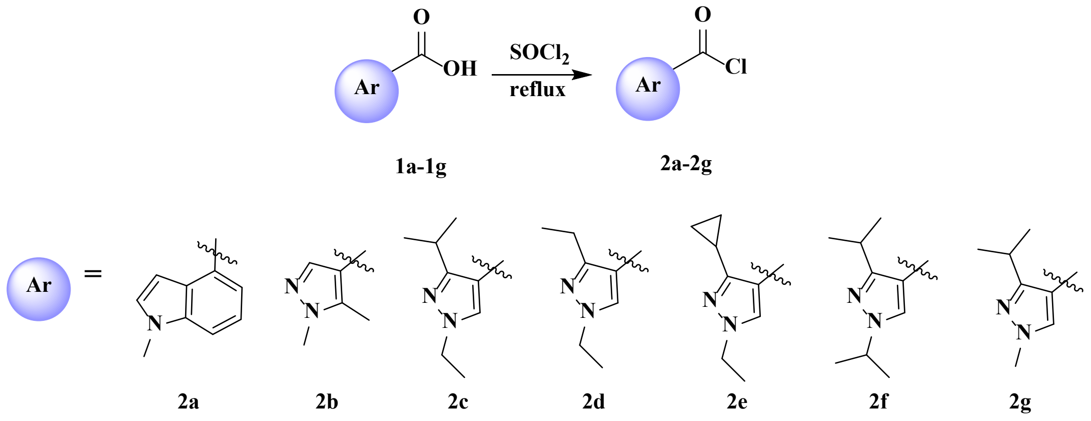


The compound **1a** (5.00 mmol) was added to dichlorosulfoxide (15 mL) and heated to reflux. The reaction was carried out for 4 h. The excess dichlorosulfoxide was removed by distillation under reduced pressure to obtain the intermediate **2a**. A similar method was used to synthesize **2b-2g**.

*2.2 Synthesis of Target Compounds**A1-A3***

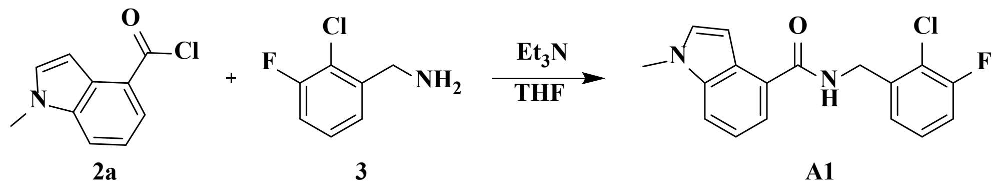


The intermediate **2a** (5.00 mmol), triethylamine ($Et_3N$, 15.00 mmol) and 2-chloro-3-fluoro-benzylamine (5.00 mmol) were added sequentially to tetrahydrofuran (THF, 15 mL) under ice bath condition. After stirring for 7 h at room temperature, the solvent was removed under reduced pressure. The crude product was purified using silica gel column chromatography with petroleum ether/ethyl acetate (V/V, 4:1) as the eluent to give the target compound **A1**.

*2.3 Synthesis of Target Compounds**A2***

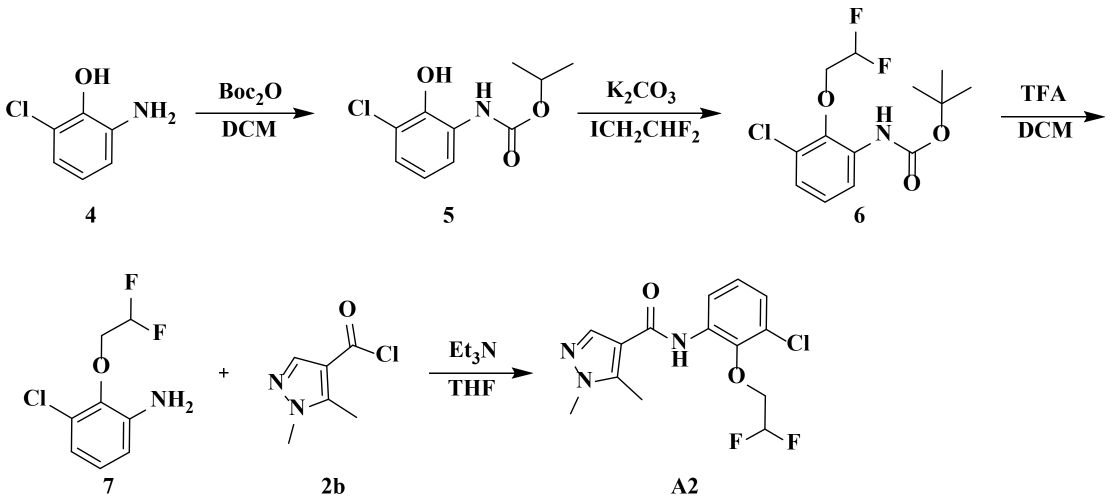


Intermediate **4** (50 mmol) and di-tert-butyl dicarbonate (25 mmol) were added to dichloromethane (DCM, 15 mL). After 12 h of reaction, excess solvent was removed under reduced pressure and separated by column chromatography to give intermediate **5**. Intermediate **5** (20 mmol) and $K_2CO_3$ (30 mmol) were added to *N,N*-dimethylformamide (DMF, 25 mL) and 1,1-difluoro-2-iodoethane (25 mmol) was added after 5 min of reaction. After 8 h of reaction, the mixture was poured into 30 mL of ethyl acetate. The organic phase was washed three times with saturated ammonium chloride solution, dried over anhydrous sodium sulfate and filtered. The excess solvent was removed under reduced pressure to give intermediate **6**. Intermediate **6** was added to 20 mL of trifluoroacetic acid (TFA)/DCM (V/V=1:3) solution. At the end of the reaction, excess solvent was removed under reduced pressure to give intermediate **7**. The intermediate **2b** (5.00 mmol), $Et_3N$ (15.00 mmol) and intermediate **7** (5.00 mmol) were added sequentially to THF (15 mL) under ice bath conditions. After stirring for 7 h at room temperature, the solvent was removed under reduced pressure. The crude product was purified using silica gel column chromatography with petroleum ether/ethyl acetate (V/V, 4:1) as the eluent to give the target compound **A2**.

*2.4 Synthesis of Target Compounds**A3**-**A7***

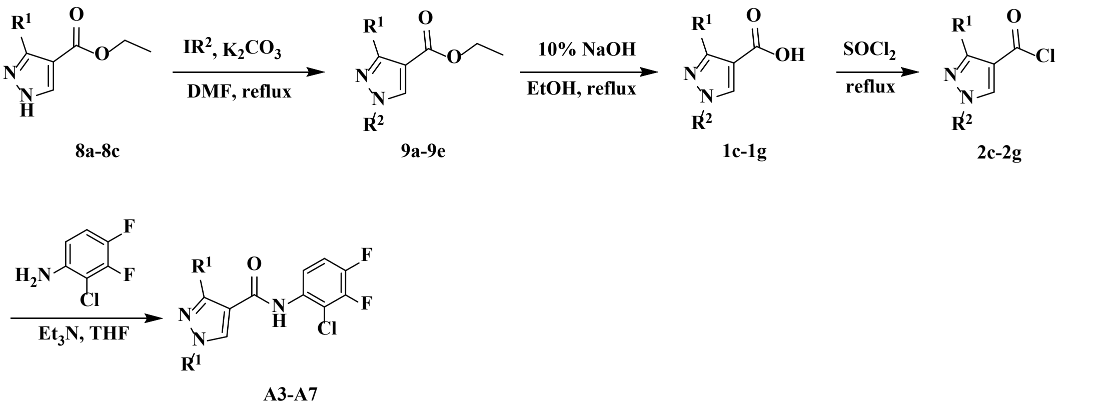


Intermediate **8a** (50.00 mmol), potassium carbonate (50.00 mmol) and 2-iodopropane were added to DMF (25 mL) sequentially. At the end of the reaction, the organic phase was washed with saturated ammonium chloride solution for 5 times, the organic phase was retained, dried over anhydrous sodium sulfate, filtered, and the excess solvent was removed under reduced pressure to give intermediate **9a**. Intermediate **9a** (30.00 mmol) and 10% sodium hydroxide solution were added to ethanol (45 mL), heated to reflux and reacted for 2 h. The reaction solution was concentrated under vacuum to remove excess ethanol, poured into ice water, adjusted the acidity with concentrated hydrochloric acid to pH=2.0, filtered and dried to obtain the intermediate **1c**. The intermediate **2c** (5.00 mmol), $Et_3N$ (15.00 mmol) and 2-chloro-3,4-difluoroaniline (5.00 mmol) were added sequentially to THF (15 mL) under ice bath conditions. After stirring for 7 h at room temperature, the solvent was removed under reduced pressure. The crude product was purified using silica gel column chromatography with petroleum ether/ethyl acetate (V/V, 2:1) as the eluent to give the target compound **A3**. A similar method was used to synthesize **A4-A7**.

*2.5 Synthesis of Target Compounds* ***A8-A21***

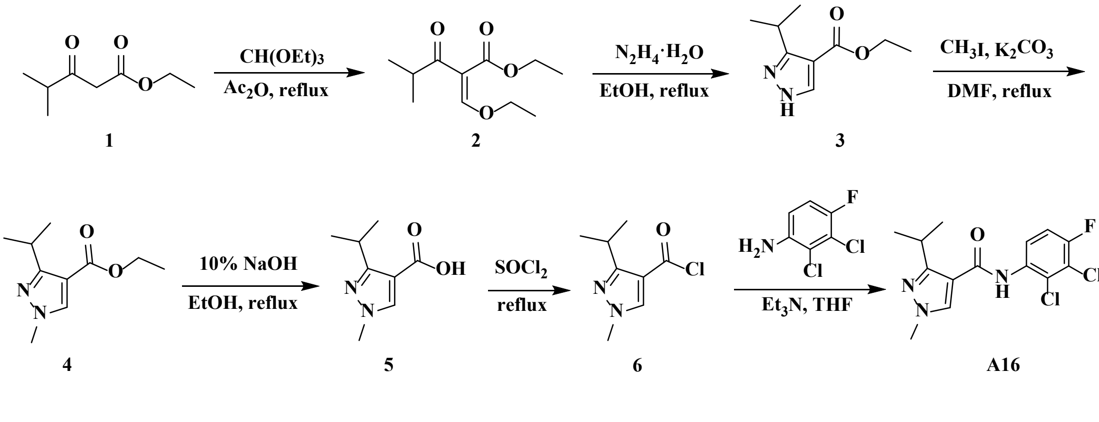

For the synthesis method of A16, please see the article. Compounds **A8-A15** and **A17-A21** have been synthesized by a similar method as an example of the synthesis of compound **A16**.

*2.6 Spectra Data of Target Compounds **A1-A21***

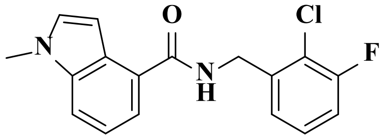

*N*-(2-chloro-3-fluorobenzyl)-1-methyl-1*H*-indole-4-carboxamide (**A1**): white solid, yield 56.13%, m.p.192–194 °C; $^{1}$H NMR (400 MHz, $CDCl_3$) $\delta$ 8.99 (t, $J$ = 5.9 Hz, 1H), 7.69 (d, $J$ = 8.3 Hz, 1H), 7.45 –7.29 (m, 5H), 7.25 (d, $J$ = 7.2 Hz, 1H), 4.58 (d, $J$ = 6.0 Hz, 2H), 3.82 (s, 3H); $^{13}$C NMR (101 MHz, $CDCl_3$) $\delta$ 168.12, 159.51, 157.04, 137.90, 135.07, 127.93–127.56 (m), 125.57–125.23 (m), 122.25, 120.15, 115.74, 115.52, 111.42, 77.25, 41.52, 30.71; HR-MS (ESI): m/z calcd for $C_{15}H_{18}ClFN_3O$ ([M+H]$^+$) 317.0852, found 317.0541.

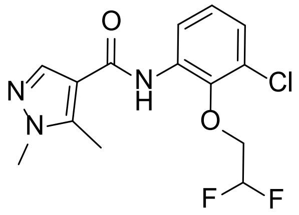

*N*-(3-chloro-2-(2,2-difluoroethoxy)phenyl)-1,5-dimethyl-1*H*-pyrazole-4-carboxamide (**A2**): white solid, yield 86.27%, m.p. 180–182 °C; $^{1}$H NMR (400 MHz, $CDCl_3$) $\delta$ 8.43 (dd, $J$ = 7.6, 2.3 Hz, 1H), 8.23 (s, 1H), 7.77 (s, 1H), 7.15–7.07 (m, 2H), 6.30–5.95 (m, 1H), 4.32 (td, $J$ = 14.3, 3.5 Hz, 2H), 3.84 (s, 3H), 2.61 (s, 3H); HR-MS (ESI): m/z calcd for $C_{15}H_{18}ClFN_3O$ ([M+H]$^+$) 330.0815, found 330.0813.

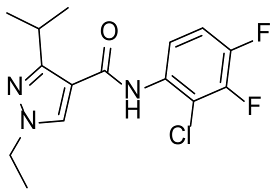

*N*-(2-chloro-3,4-difluorophenyl)-1-ethyl-3-isopropyl-1*H*-pyrazole-4-carboxamide (**A3**): white solid, yield 54.21%, m.p. 90–92 °C; $^{1}$H NMR (400 MHz, $CDCl_3$) $\delta$ 8.28 (ddd, $J$ = 9.4, 4.8, 2.4 Hz, 1H), 7.80 (s, 1H), 7.77 (s, 1H), 7.18–7.09 (m, 1H), 4.17 (q, $J$ = 7.3 Hz, 2H), 3.50 (hept, $J$ = 6.8 Hz, 1H), 1.52 (t, $J$ = 7.3 Hz, 3H), 1.37 (d, $J$ = 6.8 Hz,

6H); $^{13}$C NMR (101 MHz, $CDCl_3$) $\delta$ 161.97, 158.59, 142.44, 133.92, 109.62, 95.74, 47.36, 27.07, 21.68, 19.68, 15.13; HR-MS (ESI): m/z calcd for $C_{15}H_{17}ClF_2N_3O$ ($[M+H]^+$) 328.1023, found 328.1021.

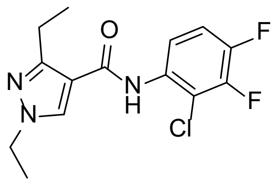

*N*-(2-chloro-3,4-difluorophenyl)-1,3-diethyl-1*H*-pyrazole-4-carboxamide (**A4**): white solid, yield 44.17%, m.p. 80–82 °C; $^1$H NMR (400 MHz, $CDCl_3$) $\delta$ 8.24 (ddd, $J$ = 9.4, 4.8, 2.4 Hz, 1H), 7.86 (s, 1H), 7.81 (s, 1H), 7.19–7.10 (m, 1H), 4.18 (q, $J$ = 7.2 Hz, 2H), 3.07 (q, $J$ = 7.6 Hz, 2H), 1.50 (t, $J$ = 7.3 Hz, 3H), 1.29 (t, $J$ = 7.6 Hz, 4H); $^{13}$C NMR (101 MHz, $CDCl_3$) $\delta$ 161.50, 147.79, 137.00, 132.08, 120.09, 115.99, 115.41 (d, $J$ = 17.2 Hz), 113.44, 44.04, 18.04, 15.60, 13.70; HR-MS (ESI): m/z calcd for $C_{14}H_{15}ClF_2N_3O$ ($[M+H]^+$) 314.0866, found 314.0866.

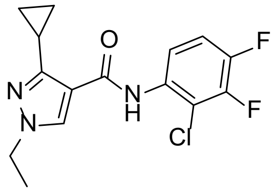

*N*-(2-chloro-3,4-difluorophenyl)-3-cyclopropyl-1-ethyl-1*H*-pyrazole-4-carboxamide (**A5**): white solid, yield 56.13%, m.p. 100–102; $^1$H NMR (400 MHz, $CDCl_3$) $\delta$ 8.48 (s, 1H), 8.27 (ddd, $J$ = 9.5, 4.9, 2.4 Hz, 1H), 7.94 (s, 1H), 7.17 – 7.09 (m, 1H), 4.10 (q, $J$ = 7.3 Hz, 2H), 2.14 (ddd, $J$ = 13.3, 8.1, 5.3 Hz, 1H), 1.47 (t, $J$ = 7.3 Hz, 3H), 1.12 – 1.02 (m, 4H).; $^{13}$C NMR (101 MHz, $CDCl_3$) $\delta$ 161.57, 132.95, 116.55, 115.36, 112.06, 77.25, 47.36, 15.20, 7.07; HR-MS (ESI): m/z calcd for $C_{15}H_{15}ClF_2N_3O$ ($[M+H]^+$) 326.0866, found 326.0870.

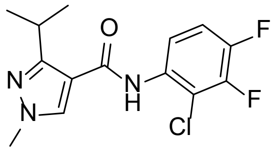

*N*-(2-chloro-3,4-difluorophenyl)-3-isopropyl-1-methyl-1*H*-pyrazole-4-carboxamide (**A6**): white solid, yield 56.13%, m.p. 148–150 °C; $^1$H NMR (400 MHz, $CDCl_3$) $\delta$ 8.26 (td, $J$ = 8.9, 5.6 Hz, 1H), 7.76 (s, 1H), 7.48 (s, 1H), 7.01–6.95 (m, 1H), 3.89 (s, 3H),

3.46 (hept, $J$ = 6.9 Hz, 1H), 1.35 (d, $J$ = 6.8 Hz, 6H); $^{13}$C NMR (101 MHz, $CDCl_3$) $\delta$ 161.54, 158.89, 131.86, 123.76, 120.07, 114.12, 111.45 (d, J = 21.0 Hz), 39.20, 27.19, 22.18; HR-MS (ESI): m/z calcd for $C_{14}H_{15}ClF_2N_3O$ ($[M+H]^+$) 314.0866, found 314.0871.

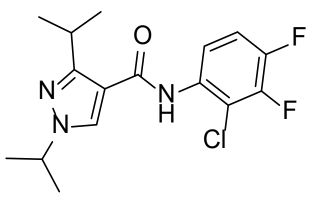

*N*-(2-chloro-3,4-difluorophenyl)-1,3-diisopropyl-1*H*-pyrazole-4-carboxamide (**A7**): white solid, yield 56.13%, m.p. 84–86 °C; $^1$H NMR (400 MHz, $CDCl_3$) $\delta$ 8.30 – 8.19 (m, 1H), 7.80 (d, $J$ = 15.4 Hz, 2H), 4.48 (hept, J = 6.7 Hz, 1H), 1.52 (d, J = 6.7 Hz, 6H), 1.36 (d, J = 6.8 Hz, 7H); $^{13}$C NMR (101 MHz, $CDCl_3$) $\delta$ 161.85, 157.53, 155.07, 147.98, 132.26, 128.18, 116.11, 115.49 (d, $J$ = 18.0 Hz), 113.83, 111.94, 77.25, 54.16, 27.48, 22.77, 22.30, 20.47; HR-MS (ESI): m/z calcd for $C_{16}H_{19}ClF_2N_3O$ ($[M+H]^+$) 342.1179, found 342.1182.

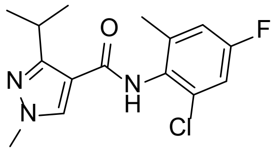

*N*-(2-chloro-4-fluoro-6-methylphenyl)-3-isopropyl-1-methyl-1*H*-pyrazole-4-carboxamide (**A8**): white solid, yield 61.24%, m.p. 130–132 °C; $^1$H NMR (400 MHz, $CDCl_3$) $\delta$ 8.33 (d, $J$ = 7.7 Hz, 1H), 7.74 (s, 2H), 7.07 (d, $J$ = 8.8 Hz, 1H), 3.90 (s, 3H), 3.53 (hept, $J$ = 6.9 Hz, 1H), 2.27 (d, $J$ = 2.0 Hz, 3H), 1.36 (d, $J$ = 6.9 Hz, 6H); $^{13}$C NMR (101 MHz, $CDCl_3$) δ 161.39, 158.80, 157.96, 131.65, 130.85, 124.66 (d, $J$ = 17.6 Hz), 123.94 (d, $J$ = 5.1 Hz), 120.09 (d, $J$ = 10.0 Hz), 115.71, 114.66, 77.26, 39.16, 27.14, 22.32; HR-MS (ESI): m/z calcd for $C_{15}H_{18}ClFN_3O$ ($[M+H]^+$) 310.1117, found 310.0997.

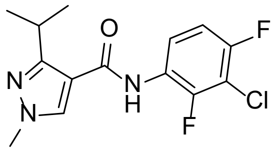

*N*-(3-chloro-2,4-difluorophenyl)-3-isopropyl-1-methyl-1*H*-pyrazole-4-carboxamide (**A9**): white solid, yield 53.14%, m.p. 102–104 °C ; $^1$H NMR (400 MHz, $CDCl_3$) $\delta$ 8.22

(ddd, $J$ = 9.5, 4.9, 2.4 Hz, 1H), 7.84 (s, 1H), 7.72 (s, 1H), 7.17–7.09 (m, 1H), 3.93 (s, 3H), 3.73 (hept, $J$ = 7.3 Hz, 1H), 1.41 (d, $J$ = 7.2 Hz, 6H); $^{13}$C NMR (101 MHz, $CDCl_3$) $\delta$ 161.74, 151.16, 137.18, 116.01, 115.50, 115.32, 114.13, 77.24, 38.23, 25.24, 19.96; HR-MS (ESI): m/z calcd for $C_{14}H_{15}ClF_2N_3O$ ($[M+H]^+$) 314.0866, found 314.0871.

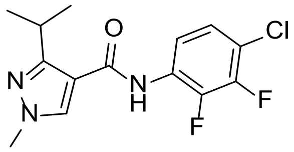


*N*-(4-chloro-2,3-difluorophenyl)-3-isopropyl-1-methyl-1*H*-pyrazole-4-carboxamide (**A10**): white solid, yield 52.71%, m.p. 148–150 °C ; $^{1}$H NMR (400 MHz, $CDCl_3$) $\delta$ 8.21–8.14 (m, 1H), 7.76 (s, 1H), 7.53 (s, 1H), 7.20–7.13 (m, 1H), 3.90 (s, 3H), 3.47 (hept, $J$ = 7.0 Hz, 1H), 1.35 (d, $J$ = 6.8 Hz, 6H); $^{13}$C NMR (101 MHz, $CDCl_3$) $\delta$ 161.35, 159.01, 127.17, 124.86 (d, $J$ = 4.1 Hz), 116.30, 114.11, 77.25, 39.20, 27.20, 22.15; HR-MS (ESI): m/z calcd for $C_{14}H_{15}ClF_2N_3O$ ($[M+H]^+$) 314.0866, found 314.0869.

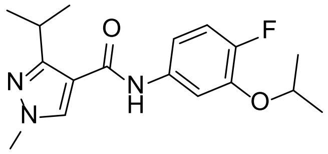


*N*-(4-fluoro-3-isopropoxyphenyl)-3-isopropyl-1-methyl-1*H*-pyrazole-4-carboxamide (**A11**): purple oil, yield 63.21%; $^{1}$H NMR (400 MHz, $CDCl_3$) $\delta$ 8.41 – 8.35 (m, 1H), 8.04 (s, 1H), 7.63 (s, 1H), 6.64 (t, $J$ = 8.7 Hz, 2H), 4.56 (hept, $J$ = 6.1 Hz, 1H), 3.91 (s, 3H), 3.76 (hept, $J$ = 7.2 Hz, 1H), 1.41 (d, $J$ = 7.2 Hz, 6H), 1.38 (d, $J$ = 6.1 Hz, 6H); $^{13}$C NMR (101 MHz, $CDCl_3$) $\delta$ 161.58, 157.65, 150.34, 147.09, 146.99, 137.18, 125.02, 120.32 (d, $J$ = 9.3 Hz), 115.15, 106.83, 106.61, 100.89, 100.62, 77.26, 71.84, 38.18, 25.10, 22.09, 20.10; HR-MS (ESI): m/z calcd for $C_{17}H_{23}ClFN_3O_2$ ($[M+H]^+$) 320.1768, found 320.1767.

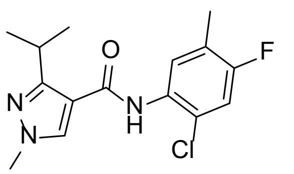


*N*-(2-chloro-4-fluoro-5-methylphenyl)-3-isopropyl-1-methyl-1*H*-pyrazole-4-carboxamide (**A12**): white solid, yield 67.15%, m.p. 148–150 °C ; $^{1}$H NMR (400 MHz, $CDCl_3$) $\delta$ 8.33 (d, $J$ = 7.7 Hz, 1H), 7.74 (s, 2H), 7.07 (d, $J$ = 8.8 Hz, 1H), 3.90 (s, 3H),

3.53 (hept, $J$ = 6.9 Hz, 1H), 2.27 (d, $J$ = 2.0 Hz, 3H), 1.36 (d, $J$ = 6.9 Hz, 6H); $^{13}$C NMR (101 MHz, $CDCl_3$) $\delta$ 8.33 (d, $J$ = 7.7 Hz, 1H), 7.74 (s, 2H), 7.07 (d, $J$ = 8.8 Hz, 1H), 3.90 (s, 3H), 3.53 (hept, $J$ = 6.9 Hz, 1H), 2.27 (d, $J$ = 2.0 Hz, 3H), 1.36 (d, $J$ = 6.9 Hz, 6H); HR-MS (ESI): m/z calcd for $C_{14}H_{15}ClF_2N_3O$ ($[M+H]^+$) 314.0866, found 314.0871.

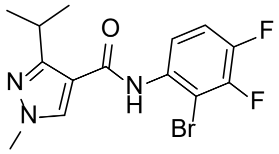

*N*-(2-bromo-3,4-difluorophenyl)-3-isopropyl-1-methyl-1*H*-pyrazole-4-carboxamide (**A13**): white solid, yield 61.47%, m.p. 150–152 °C ; $^{1}$H NMR (400 MHz, $CDCl_3$) $\delta$ 8.27 (ddd, $J$ = 9.4, 4.6, 2.4 Hz, 1H), 7.79 (s, 1H), 7.77 (s, 1H), 7.18 (q, J = 9.1 Hz, 1H), 3.91 (s, 3H), 3.60–3.48 (m, 1H), 1.36 (d, J = 7.0 Hz, 6H); $^{13}$C NMR (101 MHz, $CDCl_3$) $\delta$ 161.49, 159.00, 133.21, 131.65, 116.77–116.25 (m), 116.18, 114.33, 102.36, 77.28, 39.19, 27.15, 22.29; HR-MS (ESI): m/z calcd for $C_{14}H_{15}BrF_2N_3O$ ($[M+H]^+$) 358.0361, found 358.0364.

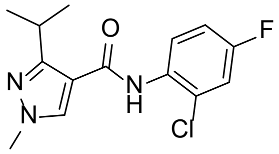

*N*-(2-chloro-4-fluorophenyl)-3-isopropyl-1-methyl-1*H*-pyrazole-4-carboxamide (**A14**): white solid, yield 60.40%, m.p. 75–77 °C ; $^{1}$H NMR (400 MHz, $CDCl_3$) $\delta$ 8.40 (dd, $J$ = 9.2, 5.7 Hz, 1H), 7.90 (s, 1H), 7.73 (s, 1H), 7.16 (dd, $J$ = 8.0, 2.9 Hz, 1H), 7.06–7.00 (m, 1H), 3.94 (s, 3H), 3.75 (hept, $J$ = 7.2 Hz, 1H), 1.43 (d, $J$ = 7.2 Hz, 6H); $^{13}$C NMR (101 MHz, $CDCl_3$) $\delta$ 161.68, 150.94, 137.20, 131.39, 123.56 (d, $J$ = 10.2 Hz), 122.78 (d, $J$ = 8.2 Hz), 116.27 (d, $J$ = 25.9 Hz), 114.61 (d, $J$ = 21.8 Hz), 114.34, 77.27, 38.19, 25.20, 19.99; HR-MS (ESI): m/z calcd for $C_{14}H_{15}ClFN_3O$ ($[M+H]^+$) 296.0960, found 296.0958.

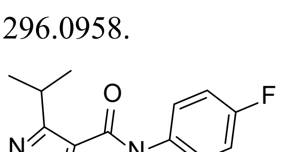

*N*-(2,4-difluorophenyl)-3-isopropyl-1-methyl-1*H*-pyrazole-4-carboxamide (**A15**): white solid, yield 65.41%, m.p. 98–100 °C; $^{1}$H NMR (400 MHz, $CDCl_3$) $\delta$ 8.30–8.21 (m, 1H), 7.68 (s, 1H), 7.57 (s, 1H), 6.87 (t, $J$ = 8.6 Hz, 2H), 3.91 (s, 3H), 3.72 (p, $J$ =

7.2 Hz, 1H), 1.39 (d, $J$ = 7.2 Hz, 6H); $^{13}$C NMR (101 MHz, $CDCl_3$) $\delta$ 161.58, 158.88, 131.73, 123.14 (d, $J$ = 8.8 Hz), 122.76 (d, $J$ = 10.4 Hz), 114.28, 111.28 (d, $J$ = 3.7 Hz), 111.06 (d, $J$ = 3.4 Hz), 103.50 (dd, $J$ = 26.7, 23.5 Hz), 77.30, 39.07, 27.11, 22.16; HR-MS (ESI): m/z calcd for $C_{14}H_{15}F_2N_3O$ ($[M+H]^+$) 280.1256, found 280.1256.

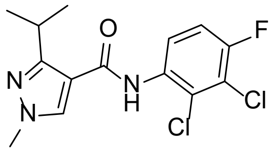


*N*-(2,3-dichloro-4-fluorophenyl)-3-isopropyl-1-methyl-1*H*-pyrazole-4-carboxamide (**A16**): white solid, yield 57.62%, m.p. 146–148 °C ; $^{1}$H NMR (400 MHz, $CDCl_3$) $\delta$ 8.36 (dd, $J$ = 9.4, 5.2 Hz, 1H), 7.91 (s, 1H), 7.72 (s, 1H), 7.15–7.09 (m, 1H), 3.93 (s, 3H), 3.72 (hept, $J$ = 7.4 Hz, 1H), 1.41 (d, $J$ = 7.2 Hz, 6H); $^{13}$C NMR (101 MHz, $CDCl_3$) $\delta$ 161.73, 151.10, 137.23, 132.55 (d, $J$ = 3.4 Hz), 123.01, 120.46, 120.14 (d, $J$ = 7.6 Hz), 114.67 (d, $J$ = 21.9 Hz), 114.13, 38.19, 25.22, 19.94; HR-MS (ESI): m/z calcd for $C_{14}H_{15}Cl_2FN_3O$ ($[M+H]^+$) 330.0570, found 330.0578.

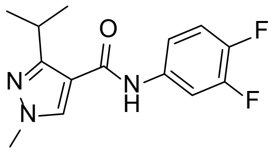


*N*-(3,4-difluorophenyl)-3-isopropyl-1-methyl-1*H*-pyrazole-4-carboxamide (**A17**): white solid, yield 68.27%, m.p. 128–130 °C; $^{1}$H NMR (400 MHz, $CDCl_3$) $\delta$ 8.34 (s, 1H), 7.65 (s, 1H), 7.57 (ddd, $J$ = 12.4, 7.2, 2.6 Hz, 1H), 7.11 (d, $J$ = 8.9 Hz, 1H), 7.03–6.95 (m, 1H), 3.81 (s, 3H), 3.68 (p, $J$ = 7.2 Hz, 1H), 1.32 (d, $J$ = 7.2 Hz, 6H); $^{13}$C NMR (101 MHz, $CDCl_3$) $\delta$ 62.39, 150.88, 137.49, 134.87, 134.75, 117.06, 116.88, 116.52–115.62 (m), 114.16, 110.08 (d, $J$ = 21.6 Hz), 38.01, 25.07, 19.90; HR-MS (ESI): m/z calcd for $C_{14}H_{15}F_2N_3O$ ($[M+H]^+$) 280.1256, found 280.1253.

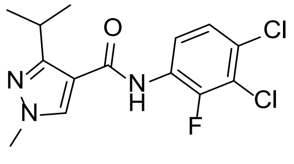


*N*-(3,4-dichloro-2-fluorophenyl)-3-isopropyl-1-methyl-1*H*-pyrazole-4-carboxamide (**A18**): white solid, yield 63.42%, m.p. 130–132 °C ; $^{1}$H NMR (400 MHz, $CDCl_3$) $\delta$ 8.24 (dd, $J$ = 9.0, 7.8 Hz, 1H), 7.68 (s, 1H), 7.66 (s, 1H), 7.25–7.20 (m, 1H), 3.91 (s,

3H), 3.69 (h, $J$ = 7.2 Hz, 1H), 1.39 (d, $J$ = 7.1 Hz, 6H); $^{13}$C NMR (101 MHz, $CDCl_3$) $\delta$ 161.74, 151.13, 137.33, 127.70, 126.51 (d, $J$ = 10.6 Hz), 125.21 (d, $J$ = 4.0 Hz), 120.14, 119.94, 113.89, 77.30, 38.18, 25.22, 19.93; HR-MS (ESI): m/z calcd for $C_{14}H_{15}Cl_2FN_3O$ ($[M+H]^+$) 330.0570, found 330.0575.

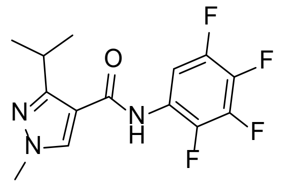

*N*-3-isopropyl-1-methyl-(2,3,4,5-tetrafluorophenyl)-1*H*-pyrazole-4-carboxamide (**A19**): white solid, yield 50.10%, m.p. 122–124 °C ; $^{1}$H NMR (400 MHz, $CDCl_3$) $\delta$ 8.24–8.14 (m, 1H), 7.68 (s, 1H), 7.56 (s, 1H), 3.93 (s, 3H), 3.70 (p, $J$ = 7.2 Hz, 1H), 1.41 (d, $J$ = 7.2 Hz, 6H); $^{13}$C NMR (101 MHz, $CDCl_3$) $\delta$ 61.63, 151.38, 137.18, 113.63, 103.84, 77.24, 38.22, 25.26, 19.90; HR-MS (ESI): m/z calcd for $C_{14}H_{14}F_4N_3O$ ($[M+H]^+$) 316.1068, found 316.1067.

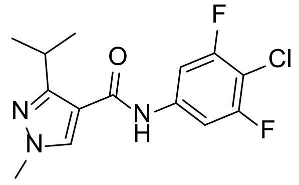

*N*-(4-chloro-3,5-difluorophenyl)-3-isopropyl-1-methyl-1*H*-pyrazole-4-carboxamide (**A20**): white solid, yield 49.76%, m.p. 152–154 °C ; $^{1}$H NMR (400 MHz, $CDCl_3$) $\delta$ 8.21 (s, 1H), 7.66 (s, 1H), 7.31 (d, $J$ = 8.6 Hz, 2H), 3.86 (s, 3H), 3.69 (hept, $J$ = 7.2 Hz, 1H), 1.36 (d, $J$ = 7.2 Hz, 6H); $^{13}$C NMR (101 MHz, $CDCl_3$) $\delta$ 162.21, 160.00, 157.59, 151.32, 138.01, 137.37, 113.92, 103.90, 103.62 (d, $J$ = 2.7 Hz), 77.29, 38.12, 25.17, 19.88; HR-MS (ESI): m/z calcd for $C_{14}H_{15}ClF_2N_3O$ ($[M+H]^+$) 314.0866, found 314.1051.

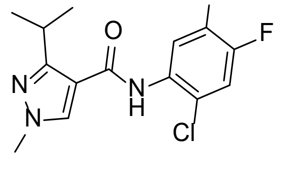

*N*-(2-chloro-4,5-difluorophenyl)-3-isopropyl-1-methyl-1*H*-pyrazole-4-carboxamide (**A21**): white solid, yield 53.84%, m.p. 134–136 °C ; $^{1}$H NMR (400 MHz, $CDCl_3$) $\delta$ 8.45 (dd, $J$ = 12.7, 8.2 Hz, 1H), 7.91 (s, 1H), 7.69 (s, 1H), 7.22 (dd, $J$ = 9.5, 7.7 Hz,

1H), 3.91 (s, 3H), 3.71 (h, $J$ = 7.1 Hz, 1H), 1.40 (d, $J$ = 7.2 Hz, 6H); $^{13}C$ NMR (101 MHz, $CDCl_3$) $\delta$ 161.50, 151.20, 137.16, 131.70, 117.56, 117.35, 116.76, 114.03, 110.45, 110.20, 77.27, 38.22, 25.22, 19.92; HR-MS (ESI): m/z calcd for $C_{14}H_{15}ClF_2N_3O$ ($[M+H]^+$) 314.0866, found 314.0869.
